# Extreme structural heterogeneity rewires glioblastoma chromosomes to sustain patient-specific transcriptional programs

**DOI:** 10.1101/2023.04.20.537702

**Authors:** Ting Xie, Adi Danieli-Mackay, Mariachiara Buccarelli, Mariano Barbieri, Ioanna Papadionysiou, Q. Giorgio D’Alessandris, Nadine Übelmesser, Omkar Suhas Vinchure, Liverana Lauretti, Giorgio Fotia, Xiaotao Wang, Lucia Ricci-Vitiani, Jay Gopalakrishnan, Roberto Pallini, Argyris Papantonis

## Abstract

Glioblastoma multiforme (GBM) encompasses brain malignancies marked by phenotypic and transcriptional heterogeneity thought to render these tumors aggressive, resistant to therapy, and inevitably recurrent. However, little is known about how the spatial organization of GBM genomes underlies this heterogeneity and its effects. Here, we compiled a cohort of 28 patient-derived glioblastoma stem cell-like lines (GSCs) known to reflect the properties of their tumor-of-origin; six of these were primary-relapse tumor pairs from the same patient. We generated and analyzed kbp-resolution chromosome conformation capture (Hi-C) data from all GSCs to systematically map >3,100 standalone and complex structural variants (SVs) and the >6,300 neoloops arising as a result. By combining Hi-C, histone modification, and gene expression data with chromatin folding simulations, we explain how the pervasive, uneven, and idiosyncratic occurrence of neoloops sustains tumor-specific transcriptional programs via the formation of new enhancer-promoter contacts. We also show how even moderately recurrent neoloops can help us infer patient-specific vulnerabilities. Together, our data provide a resource for dissecting GBM biology and heterogeneity, as well as for informing therapeutic approaches.

## Introduction

Glioblastomas (GBMs) that are wild-type for the *IDH* gene constitute the most frequent primary brain malignancy in adults (Ostrom et al., 2022). Despite their surgical resection, GBM tumors inevitably recur, are resistant to chemotherapy and highly invasive. Hence the median patient survival is ∼15 months from the time of diagnosis (Johnson and O’Neill, 2012), and therapeutic options at recurrence scarce (Wick et al., 2017; Lombardi et al., 2019). This is attributed to the documented genomic (Meyer et al., 2015; Shen et al., 2019; Kim et al., 2015a; Francis et al., 2014), epigenomic (Capper et al., 2018; Klughammer et al., 2018), and transcriptional heterogeneity of GBM tumors (Capper et al., 2018; Shen et al., 2019).

In normal tissue, the three-dimensional (3D) organization of chromosomes coordinates the activation and repression of genes to give rise to homeostatic transcriptional programs (Rada-Iglesias et al., 2018; van Steensel and Furlong 2019; Hafner and Boettinger, 2023). However, this 3D organization is disrupted at multiple levels in the context of human disease, including cancer (Spielmann et al., 2018; Ibrahim and Mundlos, 2020; Danieli and Papantonis, 2020). Structural (SVs) and copy number variants (CNVs) in tumor cells can rewire the 3D genome in ways that allow for the aberrant activation of oncogenes (Hnisz et al., 2016; Weischenfeldt et al., 2017) or the repression of tumor suppressors (Xu et al., 2022a). For example, deletion of a boundary insulating two neighboring topologically-associating domains (TADs; Beagan and Phillips-Cremins, 2020) can lead to aberrant interactions between an oncogene in one TAD and active enhancers in the other, a phenomenon known as “enhancer hijacking” (Gröschel et al., 2014; Hnisz et al., 2016; Flavahan et al., 2016; Akdemir et al., 2020a; Wang et al., 2021a). In fact, the overall distribution of somatic cancer mutations seems to be guided by 3D genome folding (Akdemir et al., 2020b).

Recently, it became apparent that by mapping 3D genome organization using Hi-C (the whole-genome variant of the chromosome conformation capture technology; reviewed in Denker and de Laat, 2016), we can simultaneously obtain a highly-resolved map of SVs and CNVs genome-wide (Harewood et al., 2017). The emergence of tools like *hicbreakfinder* (Dixon et al., 2018) and *EagleC* (Wang et al., 2022) allows for a systematic detection of SVs/CNVs in Hi-C data. Via this type of data analysis, the functional impact of SVs on subtype-specific cancer gene regulation (Xu et al., 2022a; Liu et al., 2023), as well as on a compendium of cancer lines has been investigated (Dixon et al., 2018; Wang et al., 2021a; Xu et al., 2022b). Nonetheless, our understanding of 3D genome organization in GBM remains limited due to the small number of samples analyzed to date (i.e., only 5 tumors by Harewood et al., 2017, and just 4 cell lines by Johnston et al., 2019, Wang et al., 2021b, and Yang et al., 2022).

To address this and study the impact of patient-specific SVs, we derived glioblastoma stem cell-like cells (GSCs) from 24 *IDH*-wt GBM patients—for three of which we could also sample both the primary and the relapse tumor (see **Supplementary Table 1**). It is well acknowledged that the subset of GBM tumor cells with stem-like attributes are implicated in essentially all aspects of GBM initiation, maintenance, therapy resistance, recurrence, and tissue invasion *in vivo* (Liau et al., 2017; Ricci-Vitiani et al., 2010). Given that patient-derived GSCs retain the genomic and functional traits of their tumors of origin (Lathia et al., 2015; Jacob et al., 2020; Pine et al., 2020), they hold significant potential for translational modeling of GBM. Here, we generated 28 high-resolution Hi-C datasets, and analyzed them to map structural variation in each GSC. We discovered remarkable pervasiveness and variance in SV distribution across our cohort, which gave rise to patient-specific ‘neo-TADs’ and ‘neoloops’. We combined rearranged chromosomal scaffolds with matching transcriptome and histone mark data to understand how GBM gene expression and tumor recurrence are supported by such extensive heterogeneity in their 3D genome folding.

## Results

### Pervasive structural variants cluster along GSC chromosomes

We applied *in situ* Hi-C to 28 low passage *IDH*-wt GSCs, including pairs from primary and recurrent tumors from 3 patients (**Fig. 1a**) to generate a total of ∼19 billion read pairs. Following stringent filtering, we were left with >0.4 billion valid read pairs per patient on average (63.4% mean data usage; **Supplementary Table 2**). This allowed us to produce 5-kbp resolution contact maps for each GSC, and confirm reproducibility by generating additional replicates from two randomly-selected lines (SCC>0.93; **Supplementary Fig. 1a**).

**Fig. 1.**
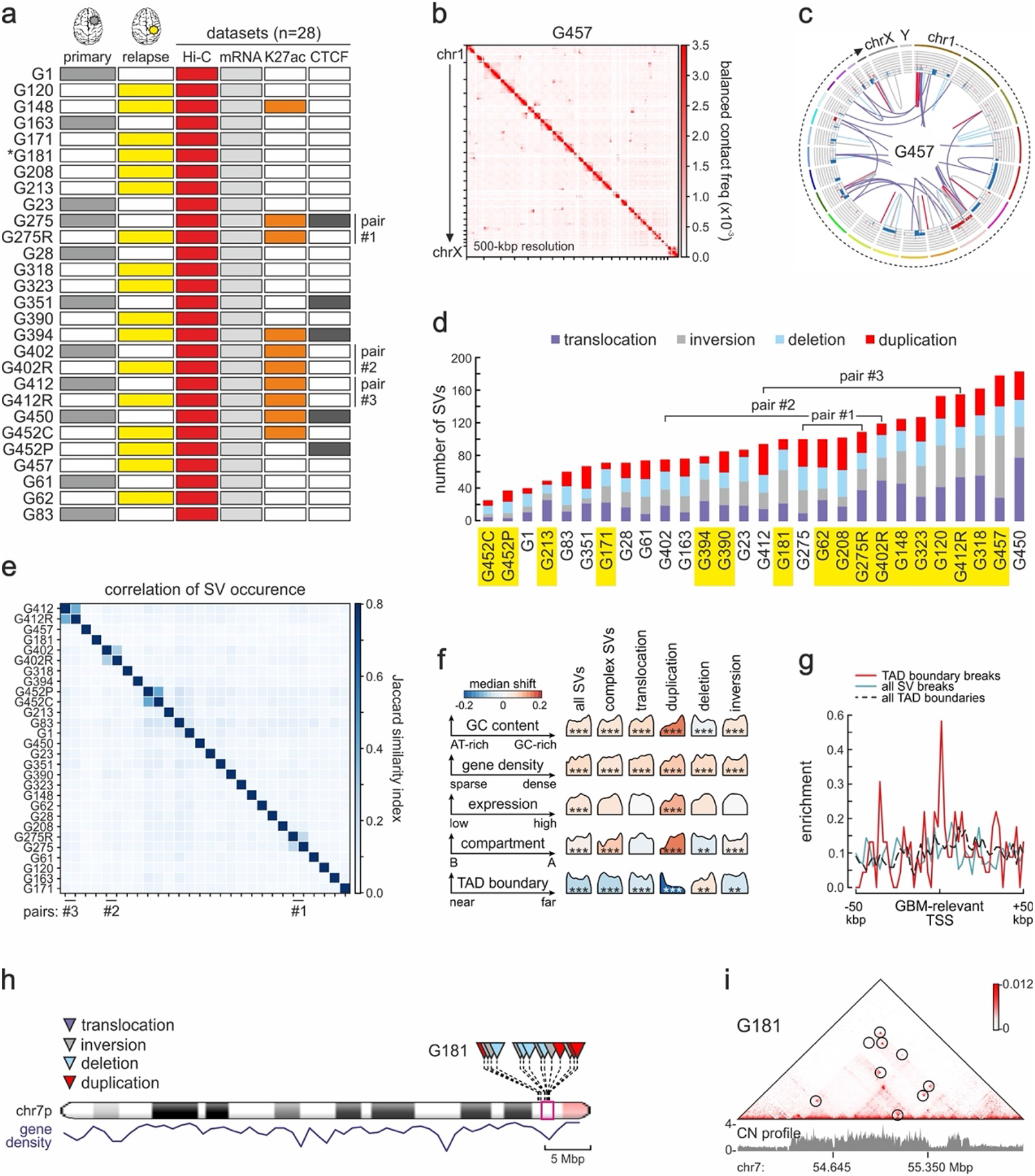
Pervasive and uneven SV occurrence discovered by Hi-C analysis of patient-derived GSCs. **a**, Overview of our cohort from 28 primary (grey) or relapse GSCs (yellow). NGS data generated from each GSC are indicated. *: WGS data is available for G181;-R designates GSCs derived from the relapse tumor in a pair, and –C/P GSCs derived from the central or peripheral part of the same tumor. **b**, Heatmap of 500 kbp-resolution Hi-C data along all G457 chromosomes. Strong interchromosomal signal represent translocations. **c**, Circos plot of SVs and CNVs detected in G457 Hi-C. Outer tracks: chromosomes; inner tracks: gain (red: >2 copies) or loss of genomic segments (blue: <2 copies); lines: translocations (purple), inversions (grey), deletions (light blue) or duplications (red). **d**, Bar plot showing the number of SV types identified in each GSC line. Lines from relapse tumors are highlighted (yellow). **e**, Jaccard similarity index of SVs discovered in different Hi-C datasets. **f**, Enrichment of breaks from all SV types (columns) relative to GC content, gene density, gene expression, A/B compartment or TAD boundaries (rows). Each density curve represents the quantile distribution of the particular genomic feature at SV breakpoints compared to random positions. **: FDR<10^-3^ or ***: FDR<10^-5^ calculated after multiple hypothesis correction on a one-sided Kolmogorov–Smirnov test based on a sample size of 5,078 genomes containing SVs. **g**, Mean enrichment of GBM-associated gene TSSs in the 100 kbp around TAD boundaries from astrocytes with an SV break in GBM (red), all SV breaks (blue) or all TAD boundaries (dashed black). **h**, Ideogram of chr7 showing SV distribution (top) and gene density (bottom) in G181. **i**, Exemplary Hi-C contact map from G181 in a 2-Mbp region of chr7 (magenta in panel h) harboring multiple SVs (circles).

We next addressed CNV prevalence in cancer cells (Shao et al., 2019) that can distort Hi-C contact maps. We verified that CNVs identified using whole-genome sequencing (WGS) data from an exemplary line, G181, were essentially identical to those computed via Hi-C data (**Supplementary Fig. 1b**). Then, we applied CNV-based matrix-balancing (Wang et al., 2021a) to Hi-C contact maps to alleviate any distortions that standard matrix balancing could not (**Supplementary Fig. 1c**). CNV-balanced matrices were next used for SV discovery in our cohort. For a comprehensive identification of SVs in our cohort, we applied *EagleC* to 5-kbp resolution Hi-C matrices (Wang et al., 2022). SVs, even those with breakpoints separated by <100 kbp, were marked by characteristic signal in our Hi-C matrices and could be detected with high sensitivity (**Supplementary Fig. 1d**). In total, we mapped 2,675 SVs across 28 Hi-C datasets, on top of 591 complex SVs (all listed in **Supplementary Table 3**). These comprised 737 (27.6%) interchromosomal translocations, plus 713 (26.7%) intrachromosomal inversions, 652 (23.4%) deletions and 573 (21.4%) duplications. Of the 1,938 intrachromosomal SVs, 57.6% were short-(<2 Mbp) and 42.4% long-range (≥2 Mbp) (for an example see **Fig. 1b,c**). Detection of SVs was robustly reproducible between replicates from the same line (mean Jaccard similarity index = 0.63; **Supplementary Fig. 1e**). As a control, *EagleC* applied to astrocyte *in situ* Hi-C (Wang et al., 2021b), only returned 7 SVs. SV occurrence across GSCs was pervasive; 16 out of 28 samples carried >80 SVs (the least number of SVs was 24 in G452C and the most was 182 for G450). Notably, relapse tumor-derived GSCs usually carried more SVs than primary ones (**Fig. 1d**). Thus, our analyses demonstrate the sensitivity and reproducibility of SV discovery in our cohort.

We next asked what the degree of SV recurrence is across our samples. Similarity analysis (**Fig. 1e**) and one-to-one comparisons of Hi-C-deduced SVs from all samples (Supplementary Fig. 2) showed remarkable heterogeneity among GSCs and little recurrence (mean Jaccard index = 0.02). Even mutations well-known to associate with GBM were only found in a subset of our samples. For example, *EGFR* locus amplification (Brennan et al., 2013) associated with SVs in 9 out of 28 lines, while *CDKN2A* deletion (Hsu et al., 2022; Funakoshi et al., 2021) was detected in 17 out of 28 lines. Finally, SVs found in GSC pairs from primary-relapse tumors of the same patient showed somewhat higher overlap (Jaccard index = 0.21). This did not increase much (Jaccard index = 0.42) even when SVs from the central and peripheral part of the same tumor (i.e., G452C/P) were considered, highlighting the intra-tumor heterogeneity of GBM.

Despite their uneven distribution across our cohort, SV breakpoint emergence correlated well with particular genomic features. For example, genomic duplications were strongly biased for strongly transcribed, GC-rich segments in the A-compartment involving breakpoints close to TAD boundaries (using astrocyte Hi-C as reference). Translocations and inversions also involved gene-/GC-rich loci, but could be both near and distal to TAD boundaries, which agrees with the notion that active gene co-association promotes rearrangements (especially translocations; Zhang et al., 2012; Sidiropoulos et al., 2022). Conversely, deletions mostly occurred in AT-rich segments of the B-compartment (**Fig. 1f**). Overall, we recorded significant enrichment for SVs occurring in the active chromatin A-compartment, particularly in gene-rich stretches, and in the vicinity of TAD boundaries (**Fig. 1f**). This is in line with the preferential occurrence of DNA double-strand break hotspots within accessible, actively transcribed chromatin (Mourad et al., 2018; Gothe et al., 2019). Notably, transcription start sites (TSSs) of genes linked to the GBM transcriptional program (as derived from DisGenet; Piñero et al., 2020) were markedly enriched at breakpoint-associated TAD boundaries (**Fig. 1g**), suggesting that TAD boundary disruption can favor oncogene dysregulation and malignant transformation (Flavahan et al., 2016; Hnisz et al., 2016; Kloetgen et al., 2020). This was also true of SVs that result in gene fusions. We identified 421 fusion events in mRNA-seq data generated from each GSC (**Fig. 1a**), but as Hi-C is more sensitive in detecting fusion positions within introns (Wang et al., 2022), we identified another 902 fusions therein (**Supplementary Table 3**) with 137 events identified by both methods. These gene fusions were expressed at significantly higher levels than their counterparts in astrocytes (**Supplementary Fig. 3a**).

Finally, rather than stochastically distributed along chromosomes, SVs show a propensity to cluster together, especially in GC-/gene-rich segments (**Fig. 1h** and **Supplementary Fig. 3b**). Such high degree of breakpoint clustering (almost 43% of SVs, i.e. 2,298 out of 5,350, were in clusters) led to complex rearrangements within relatively small (<2 Mbp) genomic stretches (**Fig. 1i** and **Supplementary Fig. 3c**). Notably, smaller chromosomes like chr12 (also remarked on in TCGA WGS data analysis; Brennan et al., 2013), 16, and 17 carried a disproportionately high density of SVs (i.e., >3.5 SVs/Mbp compared to a median of <2 SVs/Mbp; **Supplementary Fig. 3d**). As a whole, our results highlight structural variation in GSCs as a highly pervasive source of heterogeneity bound to change the 3D regulatory architecture of GBM tumors.

### GSC-specific chromatin organization blurs transcriptional subtype classification

A well-established layer of GBM heterogeneity concerns gene expression profiles. Nonetheless, analyses of bulk transcriptomic data have been used to identify three major subtypes: classical (TCGA-CL), proneural (TCGA-PN), and mesenchymal (TCGA-MES) (Verhaak et al., 2010; Neftel et al., 2019; Alhalabi et al., 2022).

To classify our GSC cohort according to these three subtypes, we used our mRNA-seq data. GSCs separated well from astrocyte profiles on PCA plots, but generated a continuum amongst themselves (**Fig. 2a**). Samples from the same patient (i.e. primary-relapse or central-peripheral GSC pairs) separated the least, suggesting that intratumor transcriptional differences are less than intertumor ones (e.g., G402 and G402R in **Fig. 2a**). We then applied single-sample gene set enrichment analysis (ssGSEA) using the CL/PN/MES signatures deduced previously (Wang et al., 2017). Out of our 28 samples, 11 showed significant enrichment (empirical *P-*value < 0.01) for CL, 9 for MES, and 2 for PN markers (**Fig. 2b**); 6 GSCs with ambiguous scores were not considered in the ensuing analysis.

**Fig. 2.**
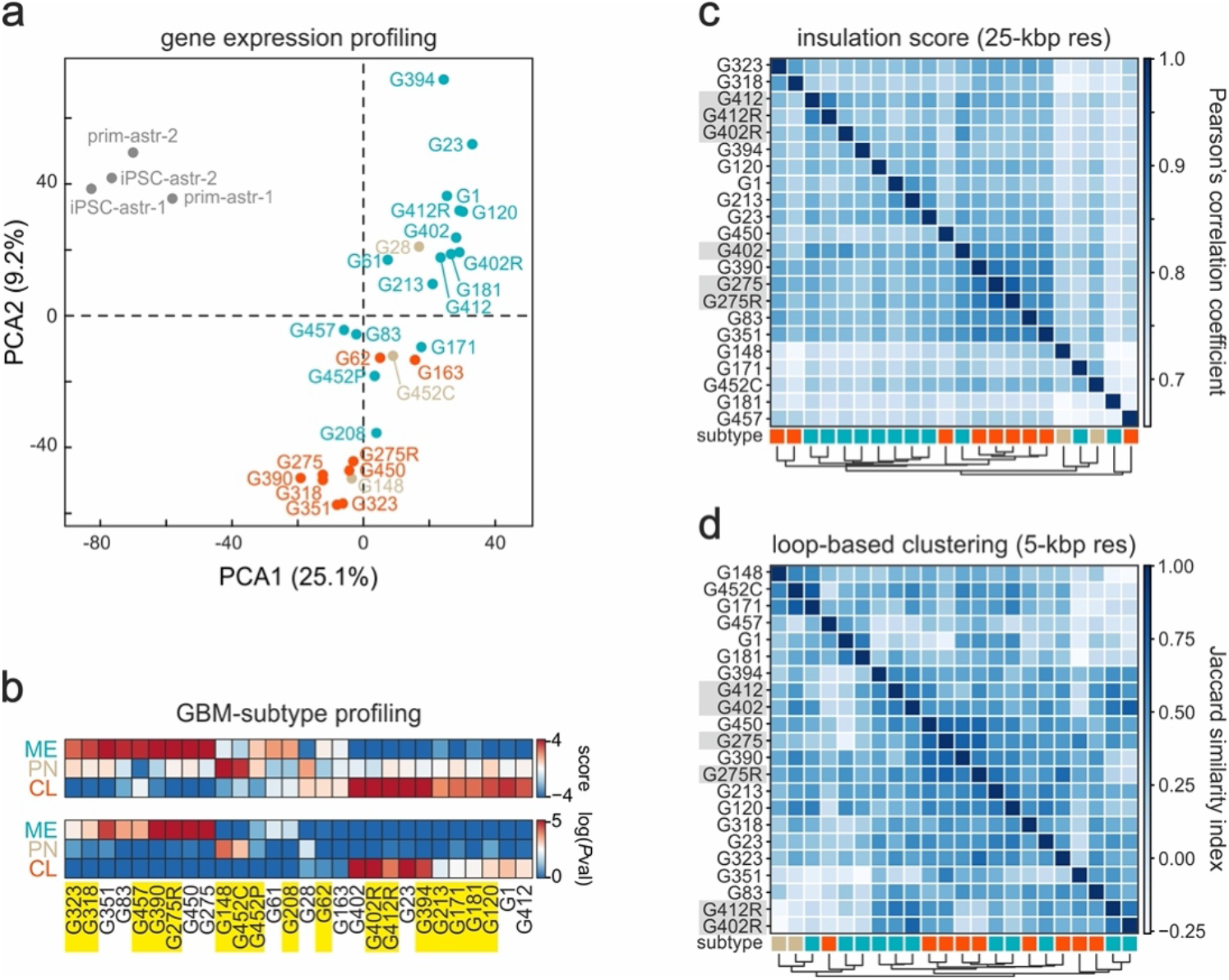
GBM transcriptional subtypes are poorly reflected in Hi-C data. **a**, PCA plot of RNA-seq replicates from 28 GSC lines classified as mesenchymal (green), classical (orange) or proneural (brown). Data from primary or iPSC-derived astrocytes provide a control. **b**, Expression-based subtyping of GSCs into mesenchymal (ME), proneural (PN) or classical (CL) based on ssGSEA enrichment scores (top) and empirically-derived *P*-values (bottom) for each signature. 22 out of 28 lines showed significant (*P*<0.01; Fisher’s exact test) association with one subtype. GSCs derived from relapse tumors are highlighted. **c**, Unsupervised hierarchical clustering of 22 GSC lines based on insulation scores calculated from 25 kbp-resolution Hi-C data. The subtype of each is indicated by colored boxes (below). **d**, As in panel c, but computing the Jaccard similarity index for loop overlap between GSCs.

Subtypes of other cancer entities (e.g., acute myeloid leukemia) were recently shown to classify on the basis of large-scale (i.e., compartmental) 3D genome organization assessed using Hi-C (Xu et al., 2022a). This motivated us to ask whether different hierarchical features in our Hi-C data would also allow classification of GSCs into the three subtypes deduced above. To this end, we used different features starting with compartments (using the first principal component of 40-kbp resolution Hi-C data eigenvectors), and continuing with insulation scores delineating TAD boundaries (calculated at 25-kbp resolution), Hi-C contacts (at 10-kbp resolution) or loops (at 5-kbp resolution). Although differential PC1 calling, reflecting GSC-specific changes in eu-/heterochromatin, broadly separated CL from MES lines (but less so PN ones; **Supplementary Fig. 4a**), all other higher-resolution features discriminated only moderately (insulation score/Hi-C contacts) or less (loops) between the subtypes (**Fig. 2c,d** and **Supplementary Fig. 4b**). We attributed this to the high structural heterogeneity that underlies individuality of each patient-derived line. Even samples with very similar transcriptional profiles like the MES lines G83 and G457 (see proximity in the PCA plot of **Fig. 2a**) share <10% of their SVs. As a result, their 3D genome features diverge profoundly and, thus, demix during clustering (see Hi-C dissimilarity in **Fig. 2c,d** and **Supplementary Fig. 4a**,b).

### GSC-specific SVs underlie neo-domains and neo-loops formation

Induction of SVs along chromosomes does not simply disturb the integrity of chromosomes and the continuity of gene loci, but also reorganize 3D spatial interactions of chromatin to give rise to new topological domains, termed ‘neoTADs’ (Franke et al., 2016; Dixon et al., 2018). We mapped neoTAD formation across all 28 Hi-Cs to identify a total of 2,222 neoTADs with a median size of 500 kbp arising from all SV types (**Supplementary Fig. 5a,b**). Different GSCs carried vastly different numbers of neoTADs (from 10 in G452C to 244 in G450; **Supplementary Fig. 5b**), which again highlights the remarkable heterogeneity of these GBM specimens. Notably, expression of genes harbored in neoTADs was consistently higher than that of genes in neighboring TADs (**Supplementary Fig. 5c**) as well as that of the same genes in astrocytes or in GSCs not forming that specific neoTAD (**Supplementary Fig. 5d**). Thus, GBM neoTADs support GSC-specific gene activity.

Similarly to neoTADs, SVs also gave rise to ‘neoloops’ characteristic of each patient-derived line (**Fig. 3a**). We used *Neoloopfinder* (Wang et al., 2021a) to identify 6,331 neoloops in locally reconstructed and normalized for allelic effects Hi-C maps from all 28 GSCs (FDR cutoff <0.95; **Supplementary Table 3**). Again, the number of neoloops in each GSC varied significantly (from 12 in G28 to 1,327 in G148, with a median of 120; **Fig. 3b**), but we saw little correlation between the number of SVs and neoloops in our cohort (ρ = 0.29).

**Fig. 3.**
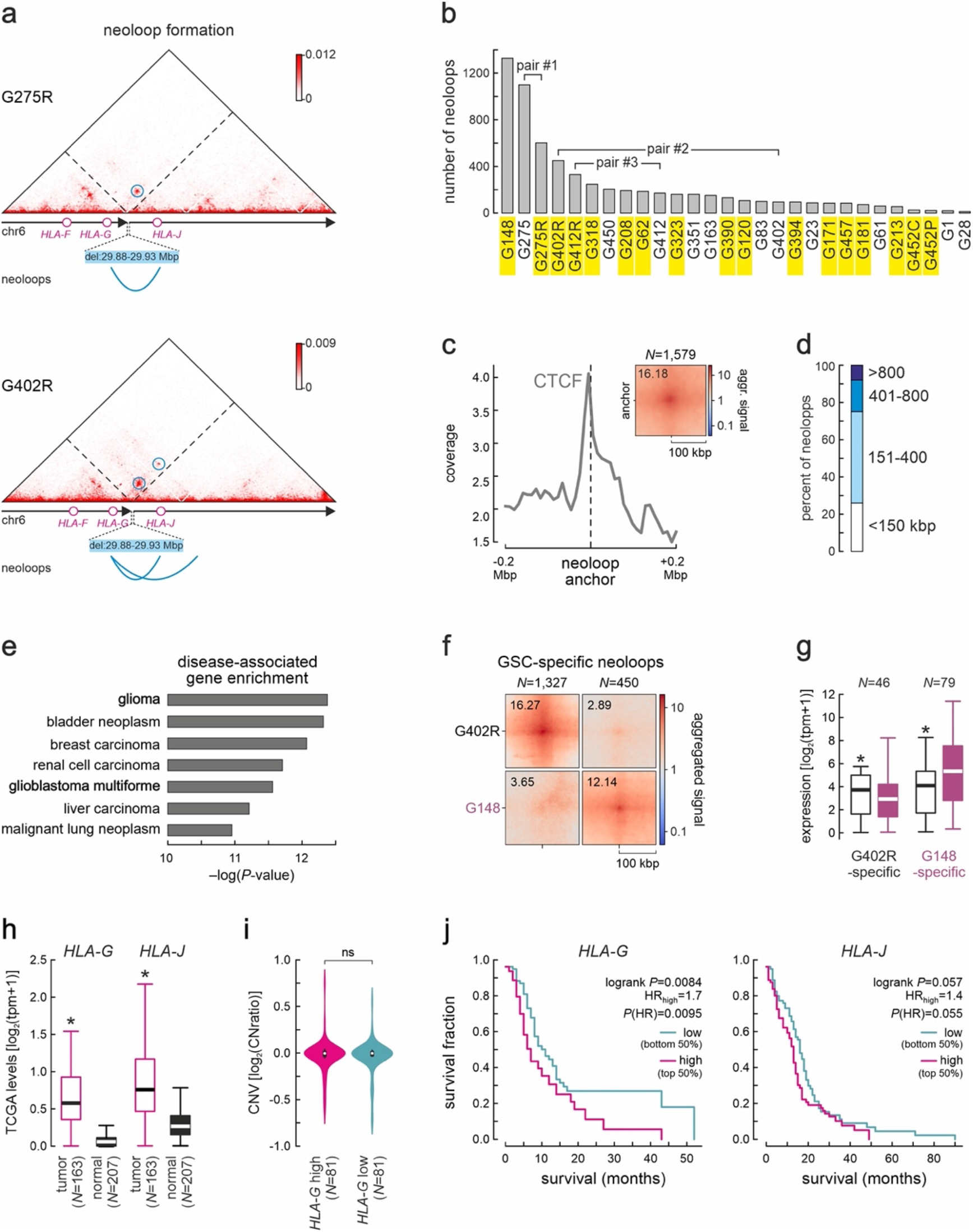
Extensive neoloop occurrence supports GSC-specific programs. **a**, Exemplary Hi-C contact maps from G275R and G402R around a 50-kbp deletion in the *HLA* locus. Neoloops forming across the breakpoint are indicated (blue circles). **b**, Bar plot showing the number of neoloops identified in each GSC line. Lines derived from relapse tumors are indicated (yellow). **c**, Line plot showing distribution of CTCF binding in the 400 kbp around all neoloop anchors. Inset: Aggregate peak analysis (APA) plot for all neoloops detected. **d**, Bar plot showing the percent of neoloops of different sizes. **e**, Signatures of neoloop-associated genes from the DisGeNET database (*P*-values calculated using Fisher’s exact tests). **f**, APA plots of neoloops specific to G402R or G148. **g**, Box plots showing expression of genes associated with GSC-specific neoloops from panel f. **h**, Box plots showing *HLA-G/-J* expression in TCGA GBM tumor and matching normal tissue. **i**, Violin plots showing copy number variation in the *HLA-G* locus from TCGA GBM tumors with high (top 50%, magenta) or low *HLA-G* expression (bottom 50%, green). *P*>0.5; two-sided Mann-Whitney U-test. **j**, Kaplan-Meier survival curves for GBM patients with *HLA-G* (left) or *HLA-J* (right) high/low expression. *P*-values were calculated using a two-sided log-rank test.

Approximately 50% of neoloops in GSCs for which we have CTCF binding information (**Fig. 1a**) were anchored at CTCF-bound sites (i.e., 772 out of 1,579; **Fig. 3c**), with 88.5% of them abiding to the expected convergent CTCF motif orientation (Rao et al., 2014). More than 90% of neoloops were <0.8 Mbp in size (**Fig. 3d**), and we identified 2,053 genes associating with neoloops (i.e., within ±5 kbp of either anchor), of which 858 were protein-coding. 131 of these protein-coding genes recurrently associated with neoloops, albeit at a low mean recurrence of 2. Amongst them, 33 (25.2%) have been reported as GBM-related (e.g., *EGFR*, *PTEN*, *MTOR*) and 29 (22.1%) as other cancer-associated genes (e.g., *AGAP2*, *SOX2*). A query for all neoloop-associated genes in DisGeNET returned a strong enrichment for genes characteristic of GBM programs (**Fig. 3e**), including genes with a high disease specificity index, like *SYF2* or *AGAP2*. Notably, neoloops sustained significantly higher expression of associated genes in a GSC-specific manner (**Fig. 3f,g** and **Supplementary Fig. 6a,b**).

Much like SV profiles that were highly heterogeneous, neoloop recurrence between GSCs was limited. For instance, the most correlated at the loop level unpaired samples, G1 and G213 (**Supplementary Fig. 4b**), shared <10% of their neoloops, while even the intra-tumor G452C/P lines shared <42%. Nevertheless, even limited recurrence becomes relevant in cases where neoloops associate with particular gene loci in different GSCs. In total, 858 protein-coding (plus 1195 non-coding) genes associated with neoloops in our cohort. Of these, 40 protein-coding (plus 59 non-coding) genes were neoloop-associated in 3 or more GSCs. On such example were the neoloops forming around the *HLA-F/-G/-J* locus following a 50-kbp deletion in 8 samples; these neoloops connected the *HLA-G* and *HLA-J* TSSs (**Fig. 3a**). *HLA-G* expression has been associated with melanoma and breast cancers (Yan et al., 2005; He et al., 2010), but only anecdotally with GBM (Wastowski et al., 2013), while *HLA-J* is considered a pseudogene with prognostic value in breast and skin tumors (Würfel et al., 2020). We analyzed TCGA data from GBM patients to find that both *HLA-G* and *-J* were significantly overexpressed in tumors versus control tissue (**Fig. 3h**), and that this was not due to gene amplification (**Fig. 3i**). Importantly, *HLA-G*/*-J* overexpression is associated with poorer patient survival overall (**Fig. 3j**). We could make similar observations for neoloops forming in the *ADAM9* locus (**Supplementary Fig. 7a**) that encodes a cell-surface protease involved in solid tumor biology (Oria et al., 2018). *ADAM9* is overexpressed in TCGA GBM tumors, not due to gene amplification, and again associates with poor patient survival (Supplementary Fig. 7b-d). This data suggests that neoloop formation is a key contributor to patient-specific gene expression, thereby helping us identify new GBM dependencies and prognostic markers.

### Enhancer-promoter neoloops explain GSC-specific gene dysregulation

Given that expression of genes was higher in GSCs where they associate with neoloops (**Supplementary Fig. 6a,b**), we wanted to further investigate how neoloops contribute to gene dysregulation in GBM. To this end, we used data from ten GSCs for which we also generated H3K27ac genome-wide profiles (**Fig. 1a**). Using this data we were able to identify putative active enhancers in each line, and saw that many of the neoloops we had charted actually represented enhancer-promoter (E-P) contacts driving GSC-specific gene expression (see the example of *IFIT1* in **Fig. 4a,b**). Of 4,343 neoloops and 11,031 non-neoloops in these ten GSC lines, E-P neoloops were significantly overrepresented (35.3%) compared to E-P non-neoloops (25.5%), while the converse applied to promoter-promoter (P-P; 9.2% neo-vs 16.9% non-neoloops) and other loops (33.3% neo-vs 66.6% non-neoloops; **Fig. 4c**). Critically, E-P neoloops linked enhancers to 192 gene TSSs in these GSCs to induce expression levels higher than those of 801 genes linked to non-neoloop enhancers. This also held true for P-P neoloops and their 243 associated genes versus 1,055 non-neoloop-associated ones (**Fig. 4d**). Similarly, the levels of 123 E-P neoloop-associated genes were significantly higher in these ten GSCs compared to lines where neoloops do not form (**Fig. 4e**). GO term analysis of E-P/P-P genes showed that they were involved in key cancer-related pathways like ‘GBM signaling’, ‘cell cycle regulation’, ‘senescence’ or ‘chromatin organization’ (**Supplementary Fig. 6c**) arguing for the tumor-specific importance of E-P neoloops.

**Fig. 4.**
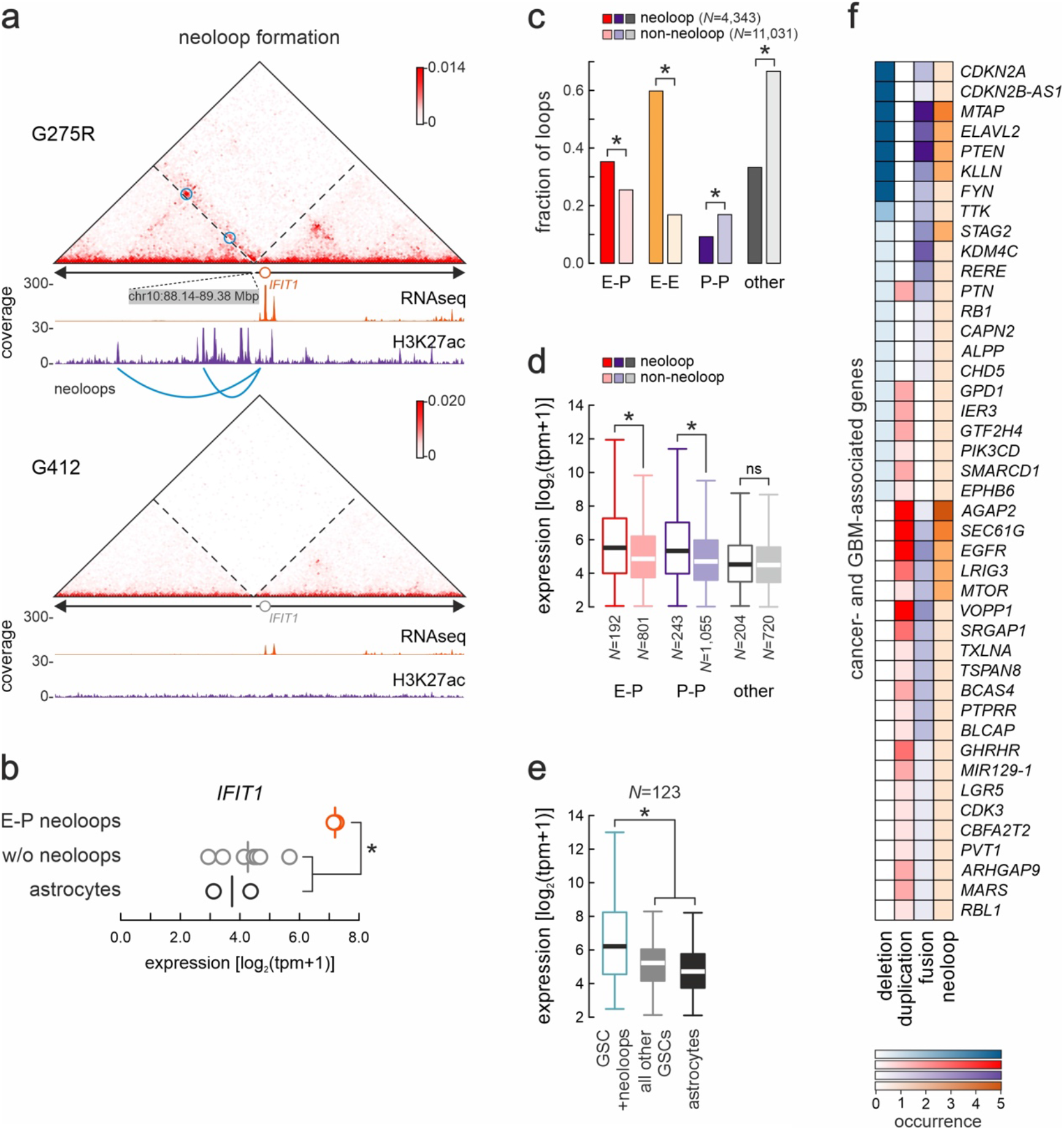
Enhancer-promoter neoloops control GSC-specific gene regulation. **a**, Exemplary Hi-C contact map from G275R around a 1.24-Mbp inversion in the *IFIT1* locus. Enhancer-promoter neoloops forming across the breakpoint are indicated (blue circles). The same locus of G412, where no inversion occurs, provides a control. **b**, Plot showing mean (line) and GSC-specific *IFIT1* RNA-seq levels (circles) in lines with (orange) or without the inversion (grey) or in astrocytes (black). *: *P*<0.01, unpaired two-tailed Student’s t-test. **c**, Bar plot showing the fraction of neo-(orange) or non-neoloops around SVs (grey) representing enhancer-promoter (E-P), enhancer-enhancer (E-E), promoter-promoter (P-P) or other interactions. *: *P*<0.01, Fisher’s exact test **d**, Box plots comparing expression of genes associated with enhancer-promoter (E-P), promoter-promoter (P-P) or other neoloop or non-neoloops. *: *P*<0.01, two-sided Mann-Whitney U-test. **e**, As in panel **d**, but for the mean expression of neoloop-associated genes in GSCs with (green) or without these neoloops (grey) or in astrocytes (black). *: *P*<0.01, two-sided Wilcoxon rank-sum test. **f**, Heatmap showing SV occurrence of known cancer-associated genes.

Finally, we compiled a list of 138 known GBM-/cancer-related genes that we could link with a neoloop and with at least one SV-linked event (i.e., with a deletion, duplication or fusion) in our cohort. Of these, 43 actually associated with at least two such events across our samples, and we saw that dysregulation could be attributed just as often to neoloop formation as to any other SV event (**Fig. 4f**). These observations suggest that E-P neoloops are a regulatory hallmark of GBM, and highly selected for in the course of tumor development to sustain favorable gene expression.

### Modeling neoloop formation underlying selective GBM dependencies

The pervasive and uneven emergence of SVs and neoloops in our cohort would inevitably give rise to GSC-specific E-P interactions and ensuing gene activation. This creates an opportunity of potential translational value: to identify tumor-specific dependencies arising from E-P neoloops activating druggable gene targets or pathways. To exemplify this, we selected G148 in which *MYC* is markedly overexpressed, not due to gene amplification, but because of a translocation between chr8 and 12. This brings into spatial vicinity the *MYC* locus (on chr8) with a cluster of enhancers (on chr12) via neoloop formation (**Fig. 5a**). This “enhancer hijacking” resulted in >10-fold increase in *MYC* levels in this one line compared to all other GSCs (or astrocytes; **Fig. 5b**), which was also reflected on MYC protein levels (**Supplementary Fig. 8a**).

**Fig. 5.**
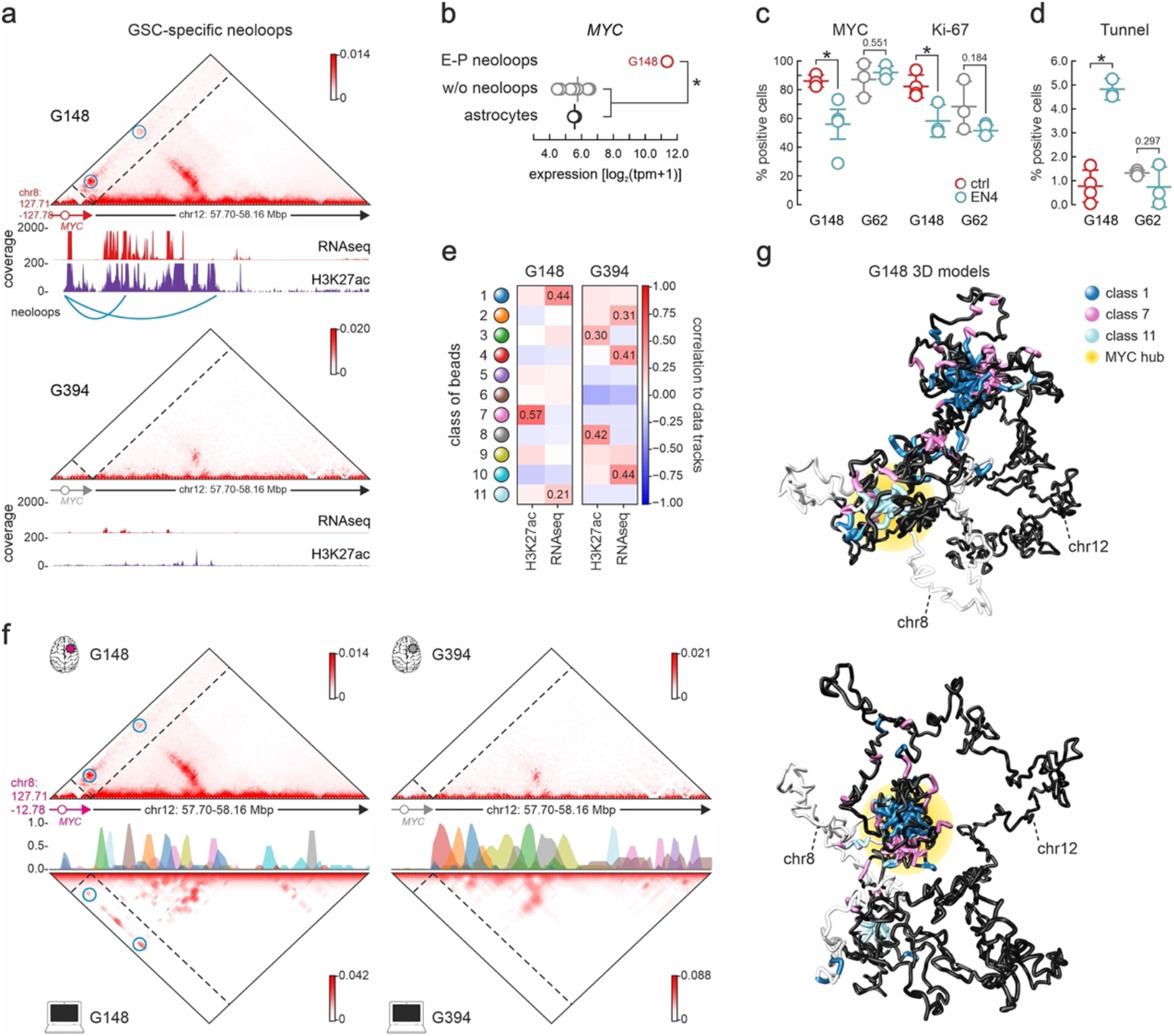
GSC-specific dependencies uncovered by neoloop analyses and simulations. **a**, Exemplary Hi-C contact map from G148 around a translocation breakpoint involving the *MYC* locus. Enhancer-promoter neoloops forming across the breakpoint are indicated (blue circles). Absence of neoloops in G394, where no translocation occurs, provides a control. **b**, Plot showing mean (line) and GSC-specific *MYC* expression (circles) in cells with (orange) or without the chr8:chr12 translocation (grey) or in astrocytes (black). *: *P*<0.01, unpaired two-tailed Student’s t-test. **c**, Plots showing the percentage of cells staining positive for MYC or Ki-67 in untreated (red) or EN4-treated *MYC*-high G148 (green) from at least 3 independent experiments; treatment of the *MYC*-low G62 provides a control. **d**, As in panel **c**, but showing the percentage of cells positive for Tunnel stainings from at least 3 independent experiments. **e**, Heatmaps showing correlation of each class of polymer beads with H3K27ac and RNA-seq data from G148 that carries the chr8:chr12 translocation (left) or G394 that does not (right). Classes with a correlation of >0.2 are shown. **f**, Left: Contact maps from Hi-C (top) or simulations (bottom) around the translocation breakpoint in G148 are shown aligned to polymer bead classes. Enhancer-promoter neoloops forming across the breakpoint are indicated (blue circles). Right: As in the left hand-side panel, but for G394 that does not carry the translocation. **g**, Representative 3D renderings of the two major configurations resulting from chr8:chr12 translocation involving the *MYC* locus. Beads from the binding classes 1, 7, and 11 that best predict folding are color-coded as in panel d, and differential *MYC*-enhancer interactions indicated (yellow halo).

We next tested whether targeting MYC would selectively inhibit G148 growth. We treated G148, as well as G62 cells, where no enhancer hijacking and no *MYC* overexpression occurs (**Fig. 5b**), with a new small molecule inhibitor, EN-4. EN-4 specifically targets Cys171 of MYC to form a covalent bond and impair its binding to target genes (Boike et al., 2021). Treatment of GSCs with EN-4 led to selective suppression of MYC and cell proliferation (marked by Ki-67expression) in *MYC*-overexpressing G148, but not in G62 (**Fig. 5c** and **Supplementary Fig. 8b,c**). Moreover, EN-4 treatment led to significant increase in DNA damage and cell death in G148 compared to G62 as deduced from Tunnel assays (**Fig. 5d** and **Supplementary Fig. 8d**).

Given the selective dependency of G148 on *MYC* overexpression, we argued that being able to predict the formation of gene-activating E-P neoloops on a patient-specific basis could guide treatment options. To achieve this, we expanded on the PRISMR *in silico* approach that was previously developed to predict ectopic interactions due to congenital disease-causing SVs (Bianco et al., 2018). In its original implementation, PRISMR could only predict ectopic interactions in *cis* by inferring binding site distribution along the polymer that best reproduces the Hi-C matrix of a genomic region in its wild-type configuration. Then, ectopic interactions are predicted by reshuffling the polymer in accordance to the SV and recalculating the new Hi-C contacts (Bianco et al., 2018). As we wanted to model a structural variant that occurs in *trans*, we modified the approach to infer binding site distribution in an extended segment of chr12 (i.e., chr12: 57.66-58.33 Mbp) using data from G275R that does not carry any SVs in the region, but in conjunction with RNA-seq and H3K27ac signal from G148 to ensure faithful Hi-C contact prediction using a probabilistic approach (see **Supplementary Fig. 9a** and Methods for details). Following optimization of binding classes (**Supplementary Fig. 9b,c**), we found that the three best-correlated ones recapitulate our input RNA-seq and H3K27ac profiles (**Fig. 5e** and **Supplementary Fig. 9a**).

In turn, the inferred classes of binding sites gave rise to highly similar contact matrices for the experiment and simulation (ρ=0.65). In G148, we could predict the formation of neoloops connecting *MYC* on chr8 to the active enhancers on chr12 (**Fig. 5f**, left), while for G394, where no translocation occurs, no neoloops were predicted (**Fig. 5f**, right). This was also reflected in 3D models rendered from the simulations, whereby a hub between the hijacked enhancers and the *MYC* locus formed in G148 (**Fig. 5g**), but not G394 data (**Supplementary Fig. 9d**). Moreover, our 3D models showed a largely mutually exclusive formation of *MYC* promoter contacts with either class-1 or-7 enhancer beads (**Fig. 5e-g** and **Supplementary Fig. 9e**). These differential conformations (**Fig. 5g**) gave rise to a heterogeneous population of 3D models, which reflected different predicted *MYC* activation levels (calculated as in Buckle et al., 2018). Models with different enhancer-promoter contacts cluster away from one another and lead to variable levels of *MYC* activation (**Supplementary Fig. 9f,g**). Thus, we can now model the impact of SVs in *cis* and in *trans* using a minimal set of epigenetic tracks to assess expression and structural heterogeneity, and potentially uncover patient-specific vulnerabilities.

### GBM relapse associates with divergence in SV occurrence and 3D genome folding

GBMs are of the most aggressively recurring tumors, with patients succumbing within ∼1 year of relapse (Ostrom et al., 2022). Our cohort included GSCs from three such primary and relapse tumor pairs (G275/R, G402/R, and G412/R; **Fig. 1a**), and we generated Hi-C, RNA-seq, and H3K27ac data from these pairs in an approach to understand their emergence.

A first observation was that, like all other GSCs in our collection, these lines also showed remarkable divergence in the number (**Fig. 1d**) and position of SVs mapped using Hi-C (**Fig. 6a** and **Supplementary Fig. 10a,b**). Relapse GSCs on average shared <35% of SVs with their primary tumor-derived counterparts (**Fig. 6b** and **Supplementary Fig. 10c,d**), while gene expression upon relapse diverged significantly in each pair (with just 30 up- and 8 downregulated genes shared by all three pairs, and <14% shared by any two; **Supplementary Fig. 10e**). Still, this extreme heterogeneity did converge to few particular pathways affected in relapse GSCs, like ‘neurogenesis’ or ‘development regulation’ (**Fig. 6c** and **Supplementary Fig. 10f**).

**Fig. 6.**
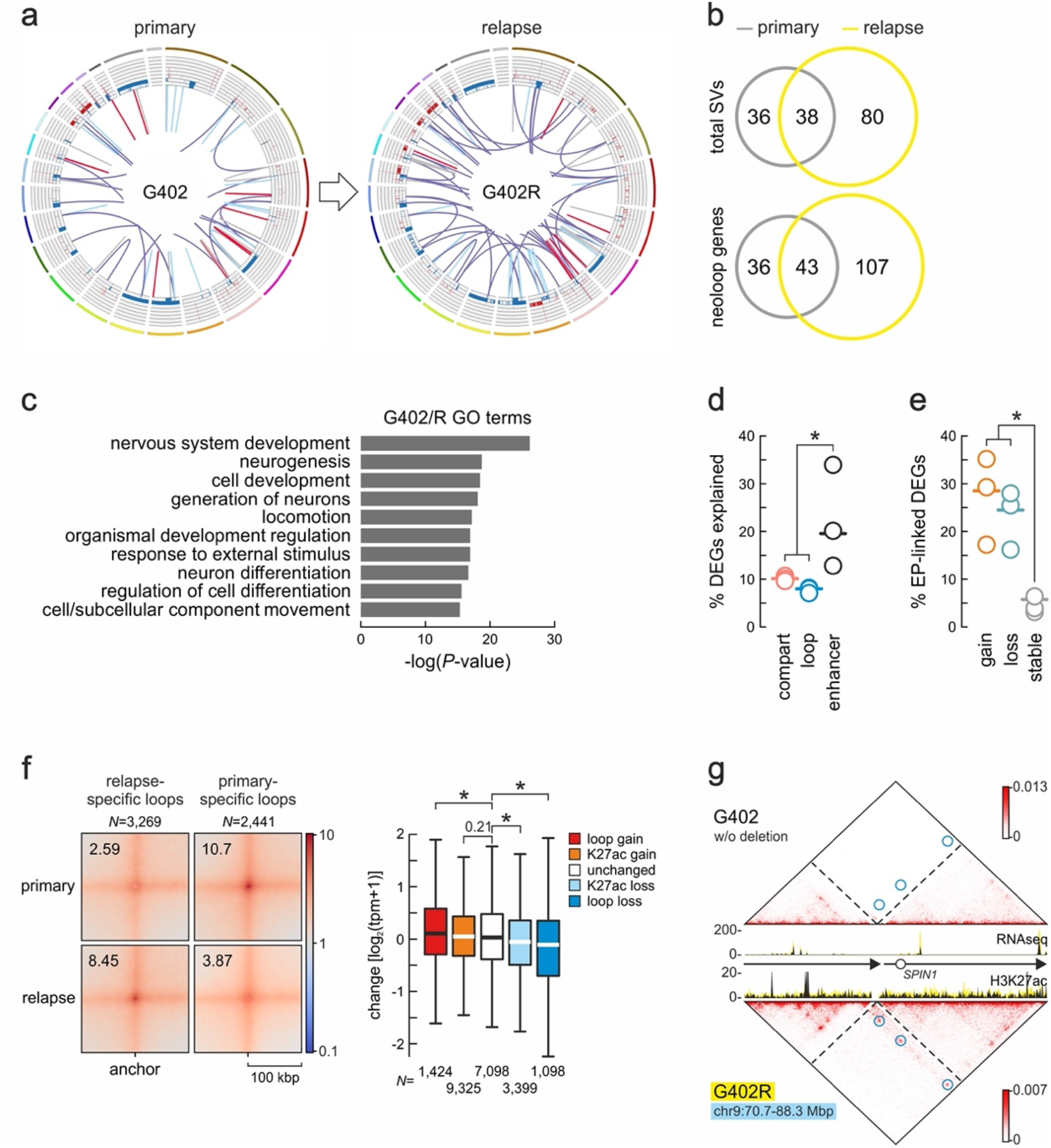
3D genome folding differentiates relapse from primary GBM samples. **a**, Circos plots of SVs and CNVs in G402 and G402R. Outer tracks: chromosomes; inner tracks: gain (red: >2 copies) or loss of genomic segments (blue: <2 copies); lines: translocations (purple), inversions (grey), deletions (light blue) or duplications (red). **b**, Venn diagrams showing shared and unique SVs (top) or neoloop-associated genes (below) in primary and relapse data. **c**, GO terms associated with differentially-expressed genes in primary versus relapse G402. **d**, Percent of differentially-expressed genes explained by A/B-compartment, loop or enhancer changes in all GSC pairs. *: *P*<0.01, unpaired two-tailed Student’s t-test. **e**, As in panel d, but for differentially-expressed genes linked to E-P loops gained (orange), lost (green) or not changed upon relapse (grey). **f**, Left: APA plot for neoloops specific to primary or relapse GSC pairs. Right: Box plots showing changes in the expression of genes associated with loops gained (red) or lost (blue), having increased (orange) or decreased H3K27ac (light blue), or not changing in relapse (white). *: *P*<0.01, Wilcoxon-Mann-Whitney test. **g**, Hi-C contact maps around a 17.6-Mbp deletion on chr9 specific to G402R shown aligned to overlaid RNA-seq and H3K27ac tracks from primary (black) and relapse samples (yellow). G402-specific neoloops are indicated (blue circles).

We next asked which level of 3D genome organization was most involved in the divergent transcriptional profiles we recorded. Switching between A- and B-compartments or association with chromatin loops each explained 10% or less of the gene expression changes seen (**Fig. 6d**). Association with enhancers though explained on average twice as many differentially-expressed genes (**Fig. 6d**), and a larger fraction of E-P loops was dynamically lost or gained between the sample pairs indicating their regulatory significance (**Fig. 6e**). On average, we identified 814 primary- and 1,090 recurrent-specific loops in the three pairs (826 and 327 for G275/R, 681 and 1,366 for G402/R, and 934 and 1,531 for G412/R; **Fig. 6f** and **Supplementary Fig. 10g,h**). We stratified these loops on whether they are specific to primary (“lost”) or relapse GSCs (“gain”), shared by a GSC pair but showing increase (“K27ac gain”) or decrease in H3K27ac signal in relapse (“K27ac loss”) or remain unchanged. After assigning genes to the P anchor of these loops, we found that expression levels of thousands of genes from all three GSC pairs were on average significantly higher (for “gain” loops) or lower (for “loss” loops) than those of genes associated with unchanged loops (**Fig. 6f** and **Supplementary Fig. 10g,h**). A considerable fraction of these loops were neoloops forming as a result of primary- or relapse-specific SVs. We looked into these and, again, relapse GSCs on average shared <41% of neoloops with their primary tumor counterparts (**Fig. 6b** and **Supplementary Fig. 10c,d**). Such neoloops often facilitated enhancer hijacking leading to aberrant gene overexpression (for an example see **Fig. 6g**). Remarkably though, we could not find any recurrence of misexpressed loci associated with neoloops or with any other SV type among the pairs. These findings suggest that GBM relapse, as reflected in GSCs, associates with a set of SVs that cannot overlap those of the primary tumor. Moreover, despite some convergence in the pathways affected, we saw marked individuality in the transcriptional programs of each GSC pair.

Finally, we used Hi-C and RNA-seq data to compare a pair of GSCs derived from the central and peripheral regions of the same GBM tumor (G452C/P). Once again, we found that the periphery shared <45% of its SVs (16 out of 36) with the center, with many SVs of G452C being lost in G452P (**Supplementary Fig. 11a,b**). This held true also for the (few) genes associated with neoloops in this pair (**Supplementary Fig. 11b**). Overall, and in line with the transcriptional divergence of these GSCs (**Fig. 2a,b**), we found pathways like “nervous system development”, ‘regulation of signaling’ or ‘ECM organization’ enriched in the central over the peripheral GSCs (**Supplementary Fig. 11c**) in conjunction with prominent GSC-specific loops in either line (**Supplementary Fig. 11d**). This data further affirms the extreme heterogeneity in GBM-derived samples, to the extent that even different parts of the same tumor diversify at the level of 3D genome architecture and regulation.

## Discussion

In this study, we generated 5 kbp-resolution Hi-C maps from 28 patient-derived GSCs and used their contact structure to identify tens to hundreds of SVs per sample. This highly resolved view of rearrangements revealed an SV distribution that was pervasive (16 out of 28 samples carried >80 SVs), yet very uneven between samples (even between GSCs derived from two different parts of the same tumor). Despite their extreme heterogeneity and largely non-recurrent nature, SVs were not stochastically distributed along chromosomes. In fact, they clustered together in hotspots correlating well with GC-/gene-rich regions preferentially located in the A chromatin compartment (i.e., transcriptionally active) and near TAD boundaries. When we focused on SV breakpoints near TSSs of genes associated with the GBM transcriptional program, we found enrichment for TAD boundaries. This suggests that disruption of such positions of 3D chromatin insulation favors oncogene activation, malignant transformation, and tumor growth (Sesé et al., 2021). Notably, in gliomas with *IDH* gain-of-function mutations, hypermethylation of CTCF sites at insulator elements that prevent binding and disrupt boundary formation (Flavahan et al., 2016). Thus, we can envisage the development of interventions that act to preserve TAD boundary integrity and counteract GBM progression in the future.

However, our cohort is exclusively *IDH*-wt and here insulation disruption is a direct result of SVs that rewire 3D chromatin folding. Still, previous “pan-cancer” analyses showed that only 14% of TAD boundary deletions actually result in a >2-fold increase in gene expression of adjacent loci (Akdemir et al., 2020a). Thus, we exploited our high-resolution Hi-C data and large number of samples assayed to focus on a key effect of GBM genomic rearrangements: the formation of hundreds to thousands (median = 120) of neoloops along each patient’s genome. Genes associated with these neoloops were not only significantly higher expressed compared to when no neoloops occur (thus, explaining intra-tumor heterogeneity), but also enriched for genes characteristic of the GBM transcriptional program. Moreover, some recurrence in neoloop-associated genes was observed (e.g., *HLA-G*/*-J* overexpression associated with neoloop formation in 8 out of 28 GSCs). Notably, a substantial fraction of these neoloops (>35%) ectopically linked gene promoters with active enhancers, which led to their activation in a tumor-specific manner. In fact, the formation of such regulatory neoloops can explain the overexpression of known GBM drivers, like *EGFR* and *MTOR*, in GSCs in cases where gene amplification or fusion does not.

We established how GBM inter-tumor heterogeneity extends to and is supported by 3D genome refolding. Then, in the absence of highly recurrent events and given the inefficacy of current treatment regimes, could 3D genomics guide personalized treatment decisions? To address this, we combined kbp-resolution mapping of 3D chromatin neo-structures with *in silico* predictions of their effects on gene expression. We expanded on the PRISMR approach by Bianco et al. (2016) to now include modeling of SV effects on 3D genome folding in *trans*. Using *MYC* overexpression via a chr8:chr12 translocation in a single GSC as an exemplar, we could show that (i) this translocation leads to the formation of two inductive enhancer-promoter neoloops; (ii) the two neoloops form in a largely mutually exclusive manner, giving rise to allele-specific conformations that explain heterogeneity in *MYC* expression; and (iii) that targeting MYC in this specific GSC with a small molecule inhibitor led to selective inhibition of its growth compared to a line not carrying the translocation and neoloops. Such patient-specific vulnerabilities may represent new opportunities for therapy, especially in the face of relapse.

GBM relapse is essentially inevitable and the major hurdle in prolonging patient survival. Studies comparing the genomic landscape of primary versus relapse *IDH*-wt glioblastomas often produce contrasting outcomes. For example, Körber et al. (2019) studied 21 primary-relapse tumor pairs using deep WGS to conclude that most of tumor evolution (incl. mutational selection) occurs even prior to primary diagnosis and, thus, relapse tumors share an overall similar landscape. This contrasts work by Kim et al. (2015b), and clinical experience, whereby GBM recurs tumultuously within a few months and relapse tumors show little genetic resemblance to primary ones. One explanation for this disparity could be the local versus distal regrowth of tumors that correlate with higher versus lower genetic resemblance (Kim et al. 2015b; Körber et al., 2019). Here, we studied three primary-relapse GSC pairs that recurred locally, but in three different brain regions (i.e., occipital, frontal, parietal). Our results on SV distribution and neoloop formation in each pair, rather argue for reduced similarity. For example, despite a consistent increase of SVs in relapse versus primary GSCs, there was an equally consistent loss of primary-specific SVs in relapse genomes. This can be explained by the two entities belonging to different (or very early diverging) tumor evolution trajectories. In addition, as our samples represent the stem cell-like compartment of GBM tumors, this could also mean that different (or even new) GSC populations emerge after resection of the primary tumor and therapy (all patients in our cohort underwent standard radiotherapy) that give rise to relapse tumors with different characteristics and resistance. We hypothesize that formation of such dynamic 3D structures as neoloops is a means for expanding regulatory options in tumor cells, and that neoloops are equally subject to tumor evolution as “classical” genomic alterations (e.g., amplifications and deletions) as they can induce significant transcriptional effects. As a result, high dissimilarity in SVs may be less telling than high dissimilarity in neoloops, as the latter can directly affect gene expression patterns. On this basis, relapse GSCs do diverge significantly from primary ones as regards their loop-level regulatory landscape despite local reemergence.

In summary, our Hi-C data constitute a valuable resource for GBM and exemplify how 3D genomics can be used to construct patient-specific chromosomal scaffolds. These can, in turn, help improve our understanding of GBM evolution and rationally identify new prognostic markers and therapeutic vulnerabilities in the face of extreme heterogeneity.

## Acknowledgements

We thank Marieke Oudelaar (MPI-NAT), Shiv K. Singh (UMG), and Kadir Akdemir (MD Anderson) for critical reading of the manuscript, and Kerstin Becker and Elisabeth Kirst from the Cologne Center for Genomics (CCG) for sequencing Hi-C libraries. This work was supported by the Italian Association for Cancer Research (AIRC; project IG2019 id. 23154 awarded to R.P.) and the Ministero della Salute (project RF-2019-12368786 awarded to R.P.), the German Research Foundation (DFG) via the Clinical Research Unit 5002 (project 426671079 awarded to A.P.), the Priority Program SPP2202 (project 422389065. awarded to A.P.), the Collaborative Research Center SFB1565 (project 469281184 awarded to A.P.) and the individual grant PA 2456/15-1 (project 455784893 awarded to A.P.). T.X. is supported by an Alexander von Humboldt Stiftung/ Bayer postdoctoral fellowship; A.D-M., I.P., and N.Ü. are members of the IMPRS Genome Science graduate school.

## Author contributions

T.X. performed all computational analyses. A.D.-M. and I.P. generated Hi-C data. N.Ü. generated CUT&Tag and RNA-seq data. M.Ba. performed simulations. M.Bu., Q.G.D., L.R.-V., G.F., and L.L. generated all GSC lines. X.W. contributed computational code. O.S.V. and J.G. performed MYC inhibition experiments. R.P. and A.P. conceived the study. T.X. and A.P. compiled the manuscript with input from all co-authors.

## Methods

### GSC generation and cell culture

GBM tumors from 24 patients who underwent surgery at diagnosis (n=11) or relapse (n=17, as 3 initially-resected patients were also part of the relapse group) at the Institute of Neurosurgery, Catholic University of Rome, were used to produce 28 GBM stem-like cell (GSC) lines. The key inclusion criteria were a GBM diagnosis (at first diagnosis or relapse; WHO Grade IV glioma), good patient functional status (Karnofsky score >70), often followed a standard Stupp protocol (for detailed profiles see **Supplementary Table 1**). Collection and processing of all samples was in compliance with the Declaration of Helsinki and approved by the Ethics Board of the hospital (Prot. ID CE 2253). Informed consent was obtained from all patients. For the generation of GSCs, surgically-removed specimens were subjected to mechanical dissociation. The resulting cell suspension was cultured in serum-free DMEM/F12 medium (ThermoFisher Scientific) containing 2 mM L-glutamine, 0.6% glucose, 9.6 mg/mL putrescine, 6.3 ng/mL progesterone, 5.2 ng/mL sodium selenite, 0.025 mg/mL insulin, 0.1 mg/mL transferrin sodium salt (Sigma Aldrich), human recombinant epidermal growth factor (hEGF; #AF-100-15, Peprotech; 20 ng/mL), basic fibroblast growth factor (b-FGF; #100-18B, Peprotech; 10 ng/mL) and heparin (2 mg/mL; Sigma Aldrich) at 37°C under 5% CO_2_. Actively proliferating cell cultures typically require 3 to 4 weeks to be established. GSCs were validated by Short Tandem Repeat (STR) DNA fingerprinting using nine highly polymorphic STR loci plus amelogenin (Cell ID™ System, Promega Inc). All GSC profiles were queried in public databases to confirm authenticity (Visconti et al., 2021). The *in vivo* tumorigenic potential of GSCs was assayed by intracranial cell injection into immunocompromised mice, resulting in tumors with the same antigen expression and histological tissue organization as the tumor of origin (Pallini et al., 2008; D’Alessandris et al., 2017).

### MYC inhibition and immunofluorescence experiments

For MYC inhibition experiments, GSCs #148 (MYC^high^) and #62 (MYC^low^) were grown as described above, but using cell culture dishes coated with growth factor-reduced Matrigel (Corning) and dissociated using Accutase (Thermo Fisher) for passaging. Once expanded, GSCs were seeded on 12 mm sterile coverslips placed in each well of a 24-well tissue culture plate. Cells were treated with either 50 µM of the small molecule inhibitor EN4 (Selleckchem; Boike et al., 2021) or with an equivalent volume of DMSO for 48 h. All experiments were performed in at least three independent biological replicates. Following drug treatment, media was aspirated and the cells fixed in 4% paraformaldehyde (PFA) for 1 h at room temperature. Cells were next permeabilized in 0.5% Triton X-100 in PBS for 10 min, and blocked with 0.5% fish gelatin in PBS for 1 h at room temperature.

MYC- and Ki67-positive cells were evaluated via immunofluorescence. In brief, primary antibody stainings were at 4°C overnight, followed by 3x 5-min PBS washes. Then, coverslips were incubated with anti-rabbit fluorophore-conjugated secondary antibodies for 1 h at room temperature. The two primary antibodies used were rabbit anti-Ki67 (Merck) and rabbit anti-MYC (Proteintech), while nuclei were also counterstained using DAPI. For Tunnel stainings, the Tunnel staining kit (Promega) was used according to the manufacturer’s instructions. Finally, coverslips were mounted and three raw images from random fields of view per coverslip were acquired using a Leica SP8 scanning confocal microscope (20x or 63x objective). Maximum intensity projection images were used and MYC mean fluorescent intensity or the percentage of Tunnel-, Ki67-, and MYC-positive cells in each sample was computed using ImageJ.

### *In situ* Hi-C and data processing

Hi-C was performed on 0.5-1 million cells from each GSC line using the Hi-C+ kit (Arima Genomics) according to the manufacturer’s instructions. Following sequencing on a NovaSeq platform (Illumina), Hi-C reads were aligned to the reference genome GRCh38 using *bwa mem* (v0.7.17) with “-SP5M”. Invalid data, including PCR duplicates and read pairs mapping to the same restriction fragment, were removed using *pairtools* (v0.3.0; Open2C et al., 2020). The *runHiC* (v0.8.4-r1; https://zenodo.org/badge/doi/10.5281/zenodo) and *cooler* (v0.8.6; Abdennur and Mirny, 2020) packages were used to construct contact matrices at various resolutions. Raw Hi-C matrices were corrected using a modified matrix balancing method to account for CNV effects and other systematic biases including mappability, GC content, and restriction enzyme sites, all processed via *Neoloopfinder* (v0.3.0.post4; Wang et al., 2021a). Stratum-adjusted correlation coefficients (SCC) between any two Hi-C contact matrices samples were calculated using Hicrep (v0.2.3) at 10-kbp resolution (Yang et al., 2017). PC1 was calculated and A-/B-compartments identified at a resolution of 50 kbp using the *cooltools* (v0.3.2; Open2C et al., 2020) *call-compartment* function. Insulation scores and TADs were identified at 25-kbp resolution using the *cooltools* (v0.3.2) insulation function. Chromatin loops were identified at 5-, 10-, and 25-kbp resolution on the basis of interaction probabilities > 0.95 and then merged using *peakachu* (v1.2.0; Samaleh et al., 2020). Significant differential loops were determined using the *diffPeakachu* function via the Gaussian mixture model of the peakachu probability score (FDR < 5%). For 5- and 10-kbp loops, we extended flanking regions by 5 kbp when searching for associated TSSs to define loop anchor genes; for 25 kbp-resolution loops no such extension was applied.

To compare chromatin organization between GSC subtypes, hierarchical structural features (PC1, insulation scores, and loops) were used for unsupervised clustering of GSC samples with significate subtype enrichment scores. For PC1 and the insulation score, pairwise correlations were calculated per each genomic bin in all samples. For loops, differential loops between GSC pairs were identified as described above, and then Jaccard similarity indexes based on shared loops were calculated, before hierarchical clustering was performed on all correlation matrices using average linkage and correlation distance metrics.

### Identification of structural variants (SVs) in Hi-C data

Structural variants, including inversions, deletions, duplications, and interchromosomal translocations, were detected and annotated using EagleC (v0.1.3; Wang, et al., 2022) on Hi-C data, which predicts SV breakpoints at single-kbp resolution and combines predictions from 5-, 10-, and 50-kbp resolutions. For 10- and 50-kbp predictions, EagleC further searches for the most probable local breakpoint coordinates within 5-kbp Hi-C contact maps so that all reported SVs are at the same resolution. In more detail, we divided the human reference genome (GRCh38) into 1-kbp bins and calculated a suite of metrics per bin to summarize a variety of properties with potential relevance to the distribution of SVs. To test for association between SV types and genome properties, each property was compared between SV breakpoint positions (randomly choosing one side of each breakpoint junction to reduce dependence between observations) and a set of 1,000 randomly-shuffled SVs, keeping the SV breakpoint ends at same distance and chromosome as those of *bona fide* ones. For each genome property and each SV type, real observations were pooled together with 1,000 sets of random ones, and rank-transformed and normalized on a 0-1 scale. Under the null hypothesis of no event-versus-property association, the ranks of the real observations would follow a uniform distribution. We tested this for each SV type using a Kolmogorov–Smirnov test with a Benjamini–Yekutieli FDR correction across the entire suite of tests, and set the threshold for significance reporting at 0.01. To define duplicated and deleted genes induced by SVs, we used both orientation information of SV breakpoints and copy number profiles. Duplications were defined as intrachromosomal SVs with −+, ++, or −− orientations, and the genomic interval between breakpoints had a copy number ratio >1.35, while deletions were also defined as intrachromosomal SVs but with the +− orientation, and the genomic interval had a copy number ratio <0.65, considering allelic heterogeneity. Copy number profiles inferred from Hi-C were used in this calculation (Wang et al., 2021a). Local Hi-C maps surrounding SV breakpoints were reconstructed and Hi-C signal across the breakpoints normalized due to the heterozygosity of the SVs and potential heterogeneity of our patient-derived GSC samples. Then, neoTADs (predicted at 25-kbp resolution) and neoloops (predicted at 5-, 10-, and 25-kbp resolutions with an FDR <0.05 and then merged) on each local reconstructed map were detected. All steps were processed using *Neoloopfinder* (v0.3.0.post4). Finally, we used RNA-seq to identify fusion genes in all GSC samples using Arriba (v2.3.0) (Uhrig et al., 2021). In parallel, we also used Hi-C processed via the EagleC (v0.1.3) *annotate-gene-fusion* function, as it can additionally detect intronic gene fusions (Wang et al., 2022). In the end, fusion genes detected via both approaches were merged to provide a final list. All SVs, CNVs, neoloops, and fusion genes are listed in **Supplementary Table 3**.

### RNA sequencing (RNA-seq) and data processing

GSCs grown to near-confluence in a T25 flask were directly lysed using Trizol (Invitrogen), total RNA was isolated using the DirectZol kit (Zymo), and used for standard poly(A)+ selection and library preparation with the TruSeq kit (Illumina). Following sequencing to at least 20 million reads on a NovaSeq platform (Illumina), reads were processed following the ENCODE pipeline (https://github.com/ENCODE-DCC/rna-seq-pipeline). Reads pairs were aligned to the human reference genome (GRCh38) and transcriptome (Gencode.v29) using STAR (v2.6.0c; Dobin et al., 2013). Gene and isoform expression quantification were performed using RSEM (v1.3.3; Li and Deqwey, 2011). Read coverage tracks (BigWig) were generated and normalized by scale factor using the *bamCoverage* function of deepTools2 (v3.5.1; Ramirez et al., 2016). Differentially-expressed genes were determined using RSEM (v1.3.3; *rsem-run-ebseq* function) with an FDR cutoff of < 0.05. For the purpose of comparing expression levels across samples, we used ”transcripts per million” (TPM) as metric. For subtype-classification. 50-gene signatures for each subtype (TCGA-PN, TCGA-CL, TCGA-MES) were used (Wang et al., 2017), and single-sample gene set enrichment analysis (ssGSEA) was conducted via R (ssGSEA). For each GSC, ssGSEA evaluated normalized enrichment scores for each signature set with TPM as input. Three two-sided *P*-values of each sample were calculated by the corresponding normalized enrichment score via the *Z2p* package and used to determine the most significant subtypes for a given GSC expression profile.

### Cleavage Under Target and tagmentation (CUT&Tag) and data processing

GSCs were lifted from plates using accutase (Sigma-Aldrich). Typically, 0.5 million cells were processed using the CUT&Tag-IT kit (Active Motif) as per manufacturer’s instructions and the resulting libraries were paired-end sequenced on a NextSeq500 platform (Illumina) to obtain at least 10^7^ reads. Read pairs were aligned to the human reference genome GRCh38 using *Bowtie2* (v2.3.4.1), PCR duplicates were removed using the *MarkDuplicates* function in Picard tools (v2.20.7), and read coverage tracks (BigWig) were generated and normalized with the RPCG parameter using the *bamCoverage* function of deepTools2 (v3.5.1; Ramirez et al., 2016). Peaks were called using SEACR (v1.3) with an FDR cutoff of <0.01 (Meers et al., 2019).

### Whole-genome sequencing (WGS) data processing

For WGS, read pairs were first mapped to GRCh38 by BWA mem (v0.7.17), and duplicate reads were removed by Picard (v2.20.7) as above. WGS-based CNV profiles and segments were calculated via the CNVkit (v0.9.9; Talevich, et al., 2016) batch function using the “--*segment-method hmm-tumor -m wgs --drop-low-coverage -- target-avg-size 25000*” parameters.

### TCGA data analysis

Kaplan-Meyer curves were generated via the Gene Expression Profiling Interactive Analysis 2 (v7.0) (GEPIA2; Tang, et al., 2019) based on 162 GBM samples from TCGA. Median gene expression values were used as a high-low group cutoff. Expression comparison between samples of glioblastoma and normal tissues were performed using GEPIA2 (v7.0) based on publicly-available TCGA and GTEx data.

### Simulations of SV impact on 3D genome folding

In order to predict neoloops forming as a result of translocations, we used a polymer physics based approach previously used to predict ectopic interactions in cis arising in congenital disease-causing structural variations, PRISMR (Bianco, et al., 2018). PRISMR models chromatin as a polymeric structure bearing sites of potential binding by proteins represented as floating particles in solution (Barbieri et al., 2012; Chiariello et al., 2016). The thermodynamic properties of this model can be used to infer the binding site distribution along the polymer that best reproduces the Hi-C matrix of the genomic region lacking the structural variation, while ectopic interactions are predicted by reshuffling the polymer in accordance with the variation and recalculating the new Hi-C contacts. Here, we modified the PRISMR approach to simulate neoloop formation by a translocation involving the *MYC* locus on chr8 and a large intergenic segment of chr12 (chr8: 127.71-127.78 Mbp; chr12: 57.7-58.155 Mbp); this occurs only in sample G148 of our cohort. Similar data from G394, where the translocation does not occur and *MYC* is activated provided a negative control.

To predict the best binding sites distribution for the hybrid region in G148, we first inferred them on an extended region on chr12 (i.e., chr12: 57.66-58.33Mbp) using Hi-C data from G275R that carries no SVs across this segment. We then correlated the binding site distribution deduced from G275R with RNA-seq and H3K27ac CUT&Tag data from the same region in G148 to ensure good prediction of Hi-C contacts via a probabilistic approach using these correlation to extend binding site distribution prediction around the *MYC* locus on chr8 (i.e., chr8: 127.71-127.78 Mbp). Our approach repurposed PRISMR that finds the best minimum of the difference between the real Hi-C matrix and the reconstructed Hi-C matrix via a simulated annealing (SA) optimization procedure spanning the space of binding site distributions for a given number of classes. A contact matrix is then reconstructed via a mean field approximation using contact probability profiles characterized in the standard coil-globule theory of polymer physics (DeGennes, 1979; Barbieri et al., 2012; Bianco et al., 2018). Estimation of the best number of binding sites classes and the best λ (i.e., the regularization term used in PRISMR SA to penalize total binding site abundance and reduce overfitting) was as previously described (Bianco et al., 2018). In brief, SA was executed for a range of λ values, and the best λ was selected when the cost function raised ∼10% above the starting plateau (**Supplementary Fig. 8b**). Similarly, SA was executed for an increasing number of binding sites classes, *M*, until the cost function did not show significant reduction (*M=11* was selected; **Supplementary Fig. 8c**). For this, experimental (input) Hi-C data was first smoothed via a Gaussian filter (0.5 bin), and similarity between the simulated and original contact matrices was estimated using distance-corrected Pearson’s correlations (which were 0.65 and 0.45 for G148 and G394, respectively). Also, to account for chromatin persistence length effects in our 5-kbp resolution deduced Hi-C matrices, SA was applied independently for different monomer lengths by interpolating and scaling contact probability profiles accordingly, albeit with a significant increase in computing burden. To speed up optimization convergence, we modified SA to generate, at every iteration, multiple tentative modifications of the binding site configuration (rather than one in Bianco et al., 2018) that were simultaneously evaluated. This allowed us to estimate an optimal monomer length ∼20% longer than the 5kb resolution.

To extend prediction of binding sites distribution to the *MYC* locus on chr8, we applied a probabilistic approach, using RNA-seq and H3K27ac data as a bridge between chromosomes. If P_C_ is the probability to find the binding sites class C in the region of interest, P_T_ is the probability to find the epigenetic track T, and *corr(C,T)* is the correlation between C and T, then the conditional probability *P(C|T)* to observe C given T can be obtained by inverting the following equations:

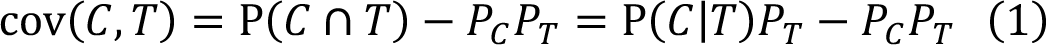

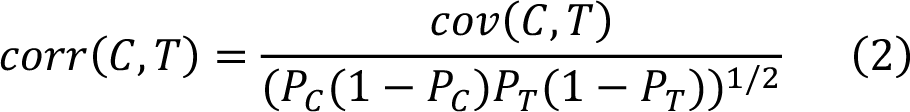

Here we define *corr(C,T)* as the correlation between the PRISMR-inferred best binding sites distributions (*C*) and the RNA-seq and H3K27ac tracks on the extended region chr12:57.66-58.33Mbp (*T*), and *P_C_ (P_T_)* as the frequency of observing C (T) in the same region. Once we have *P(C|T)*, we can estimate the probability to find a binding site of class C in position x in region chr8:127.71-127.78Mbp from the frequency of T as follows:

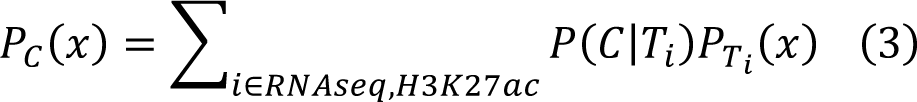

In this formula we neglected the intersection terms between *P_RNAseq_* and *P_H3K27ac_* as their correlation is quite low (<0.2). When applying (3) we considered only *(C,T)* couples with a correlation of >0.2. Equations (1-2) follow from the very definition of correlation and covariance where *cov(C,T) = cov(1_C_,1_T_)* and *1_X_* is the indicator function, so the expected values in the covariance equal the probabilities:

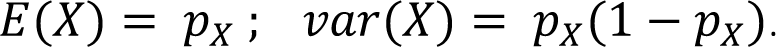

Finally, to predict the three-dimensional structure and dynamics of the genomic region bearing the translocation, we employed the SBS model via Molecular Dynamics simulations in a classical Langevin and velocity-Verlet framework with standard parameters (Barbieri et al., 2012; Chiariello et al., 2016; Bianco et al., 2018). The energy of interaction between binding sites and binders was set to 4 K_B_T, while the binders’ concentration was set to 100 nmol/liter. Randomly generated polymers and binder configurations were allowed to evolve and find the steady state before measuring the probability of contact. From the SBS predicted structures we estimated the degree of *MYC* triplet colocalization with region A and B with respect to what expected by random independent pair-wise probability via the correlation coefficient:

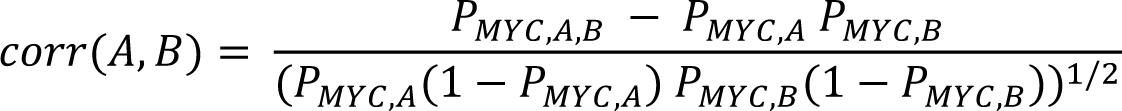

From the SBS polymer distance matrix we also estimated the level of in-silico single-allele MYC expression with respect to the average level F following the formula in Buckle, et al., 2018:

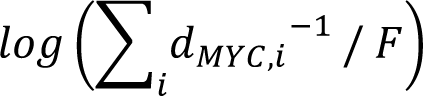

where *d_MYC,I_* is the distance between *MYC* and *i* corresponding to a H3K27ac or RNA-seq peak.

### Statistics and reproducibility

All *P*-values were calculated using R, and their results considered significant if *P*<0.01, unless stated otherwise.

## Data availability

Due to patient protection policies, raw Hi-C, mRNA-seq, and H3K27ac/CTCF CUT&Tag data may only be released upon request and ethics approval. Processed data that do not contain identifiable information can be accessed via the NCBI Gene Expression Omnibus (GEO) under accession number GSE229966. GBM-associated genes were obtained from the DisGenet Database (v7.0) (Piñero et al., 2020). A list of cancer-related was sourced from: http://www.bushmanlab.org/assets/doc/allOnco_May2018.tsv, and gene-level copy numbers of TCGA samples from the cBioPortal (https://www.cbioportal.org/). Astrocyte RNA-seq data was sourced from Santos et al., 2017.

## Code availability

All code used to analyse Hi-C, RNA-seq, WGS and CUT&Tag data is available at https://github.com/xieting0603; the custom code used to perform simulations is available at https://github.com/marianoimperatore/MeanFieldChromatin.git.

**Supplementary information** accompanying this manuscript includes Supplementary Figs 1-11 and Supplementary Tables 1-3.

## SUPPLEMENTARY INFORMATION

**Supplementary Fig. 1.**
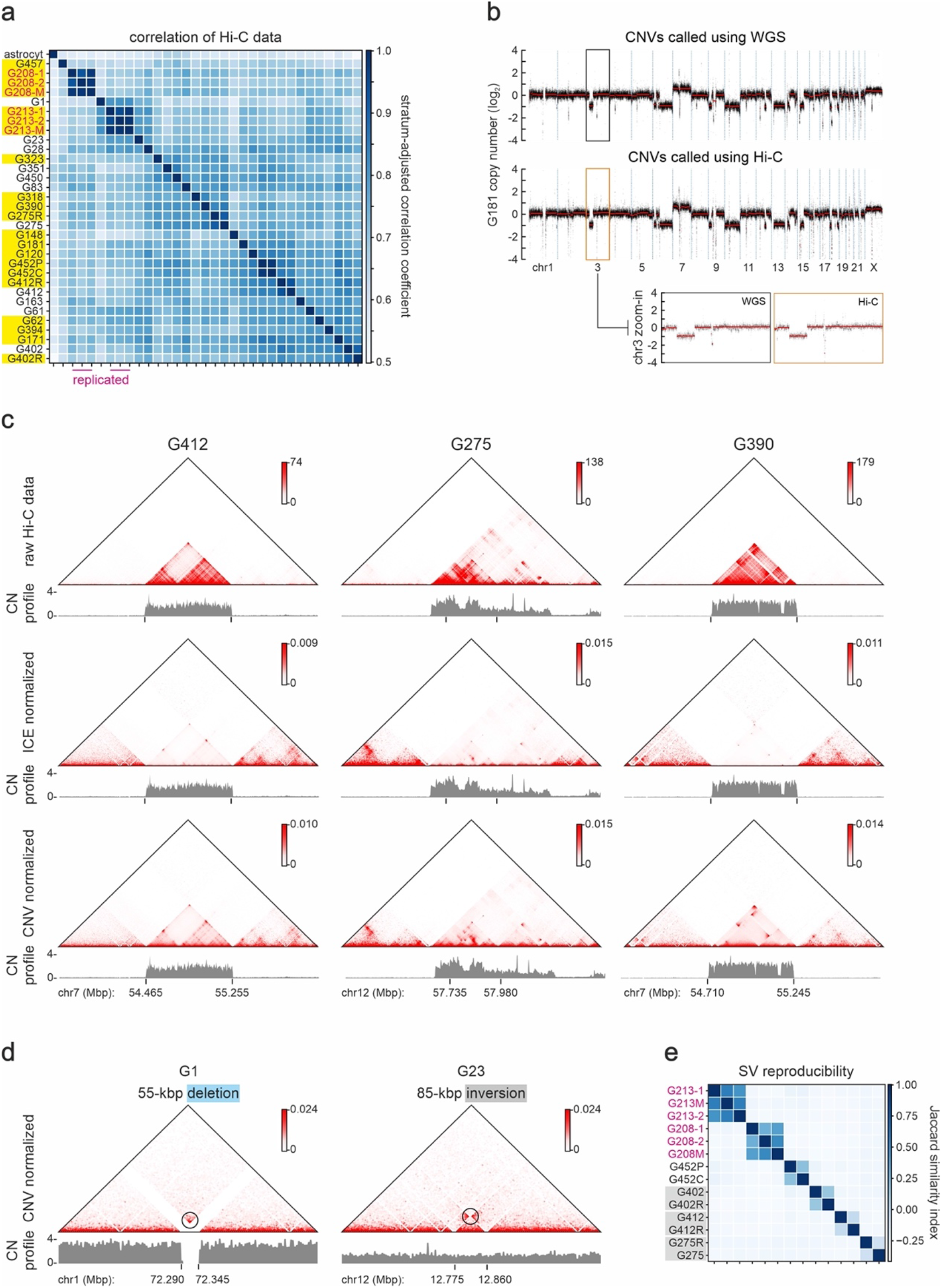
Evaluation of Hi-C reproducibility, CNV segmentation, and normalization. **a**, Heatmap showing stratum-adjusted correlation between 10 kbp-resolution raw Hi-C contact matrices from all GSC lines. Biological replicates from the same line (−1/-2) and their merged map (M) are indicated. Data from relapse GSCs are highlighted (yellow). **b**, Comparison of whole-genome CNV computations using WGS (via CNVkit) or Hi-C data from G181 (via Neoloopfinder). A zoom-in for the CNVs identified along chr3 is provided. **c**, Comparison of 5 kbp-resolution raw (top), ICE-normalized (middle) or CNV-normalized Hi-C contact matrices (bottom) around exemplary amplified regions (CNV profiles aligned below). **d**, Exemplary 5-kbp resolution Hi-C contact maps showing signal characteristic of short-range SVs for a deletion in G1 (left) and an inversion in G23 (right). **e**, Heatmap showing similarity of SVs discovered in Hi-C data of 12 exemplary GSCs. Biological replicates from the same line (−1/-2) and their merged map (M) are indicated.

**Supplementary Fig. 2.**
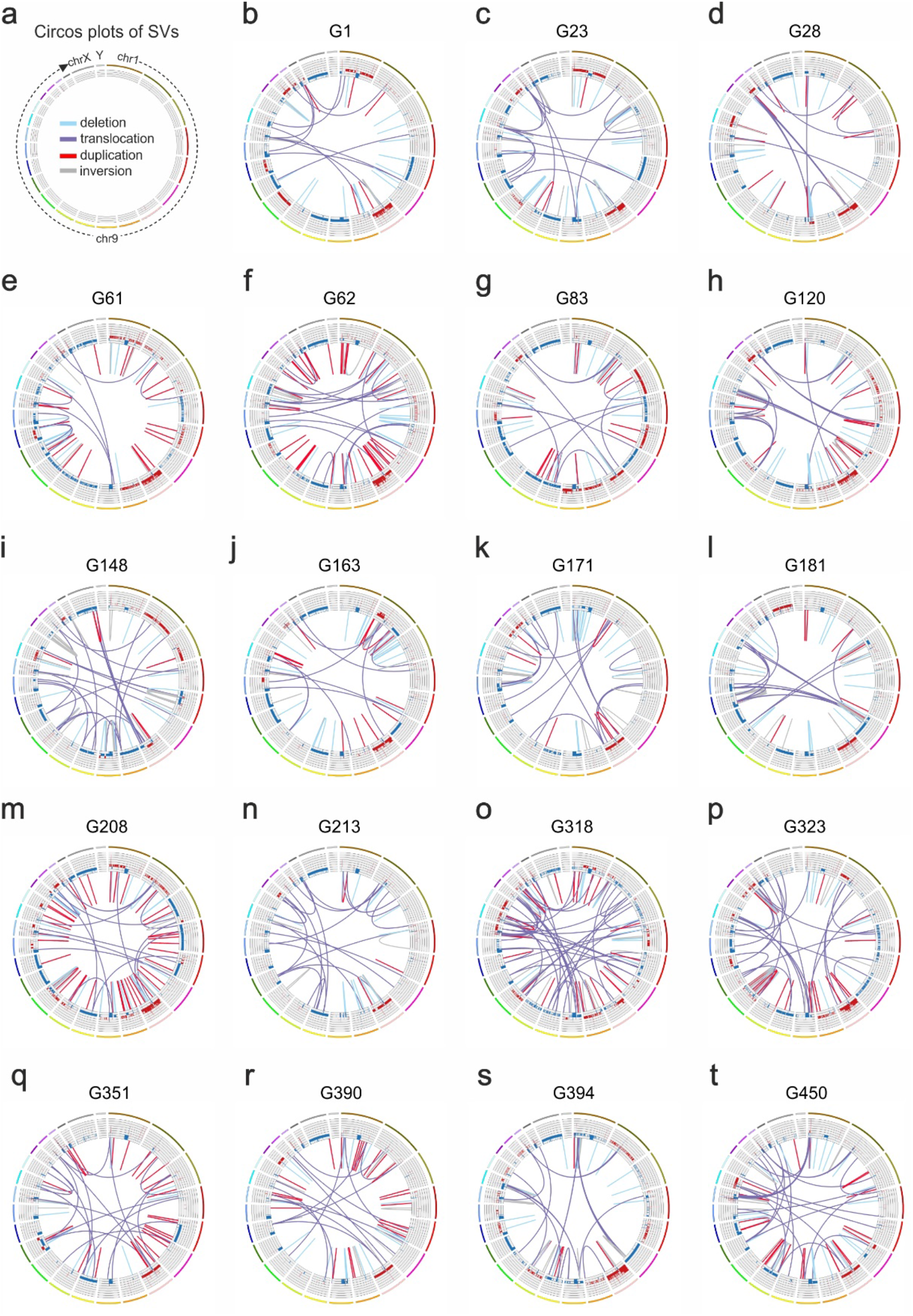
Distribution of SVs across GSC lines. **a**, Key showing the positions of chromosomes (outer tracks) and the color code for SVs in Circos plots (inversions – grey; deletions – light blue; duplications – red; translocations – purple). **b-t**, Circos plots of SVs and CNVs detected in 19GSC Hi-C datasets. Inner tracks: gain (red, >2 copies) or loss of genomic segments (blue, <2 copies); lines: SVs as detailed in panel a.

**Supplementary Fig. 3.**
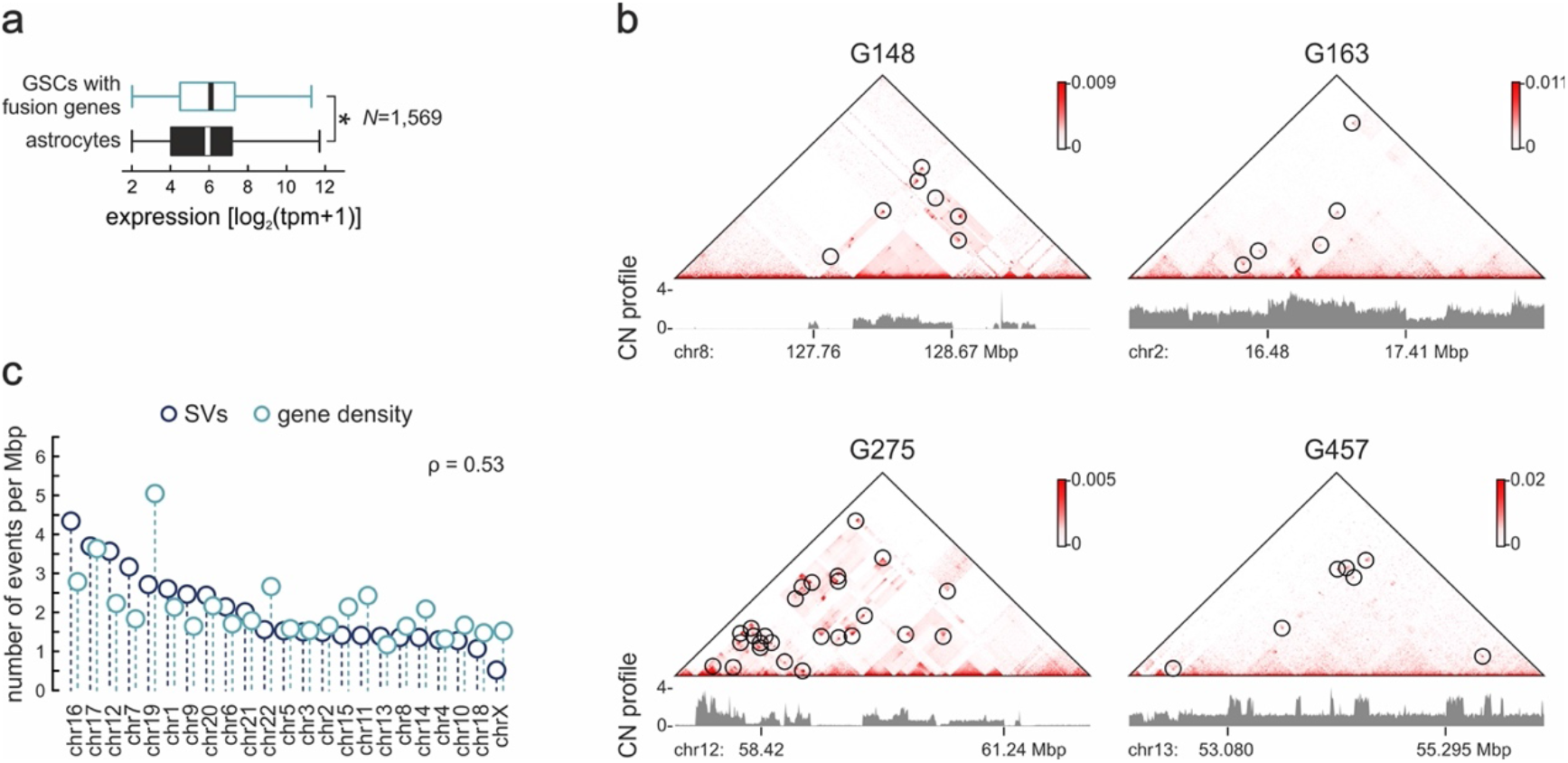
Clustered SV occurrence along GSC chromosomes. **a**, Box plots showing expression levels of 1,569 gene fusions in GSCs (green) compared to their individual counterparts in astrocytes (black). \**P* <0.01, two-sided Wilcoxon rank-sum test. **b**, Exemplary Hi-C contact maps from 4 GSC lines showing SV clustering in 3-Mbp stretches of different chromosomes. **c**, Lollipop plots showing the number of SVs (dark blue) or genes per Mbp of each chromosome (light blue). Pearson’s correlation coefficient (ρ) for the two datasets is calculated.

**Supplementary Fig. 4.**
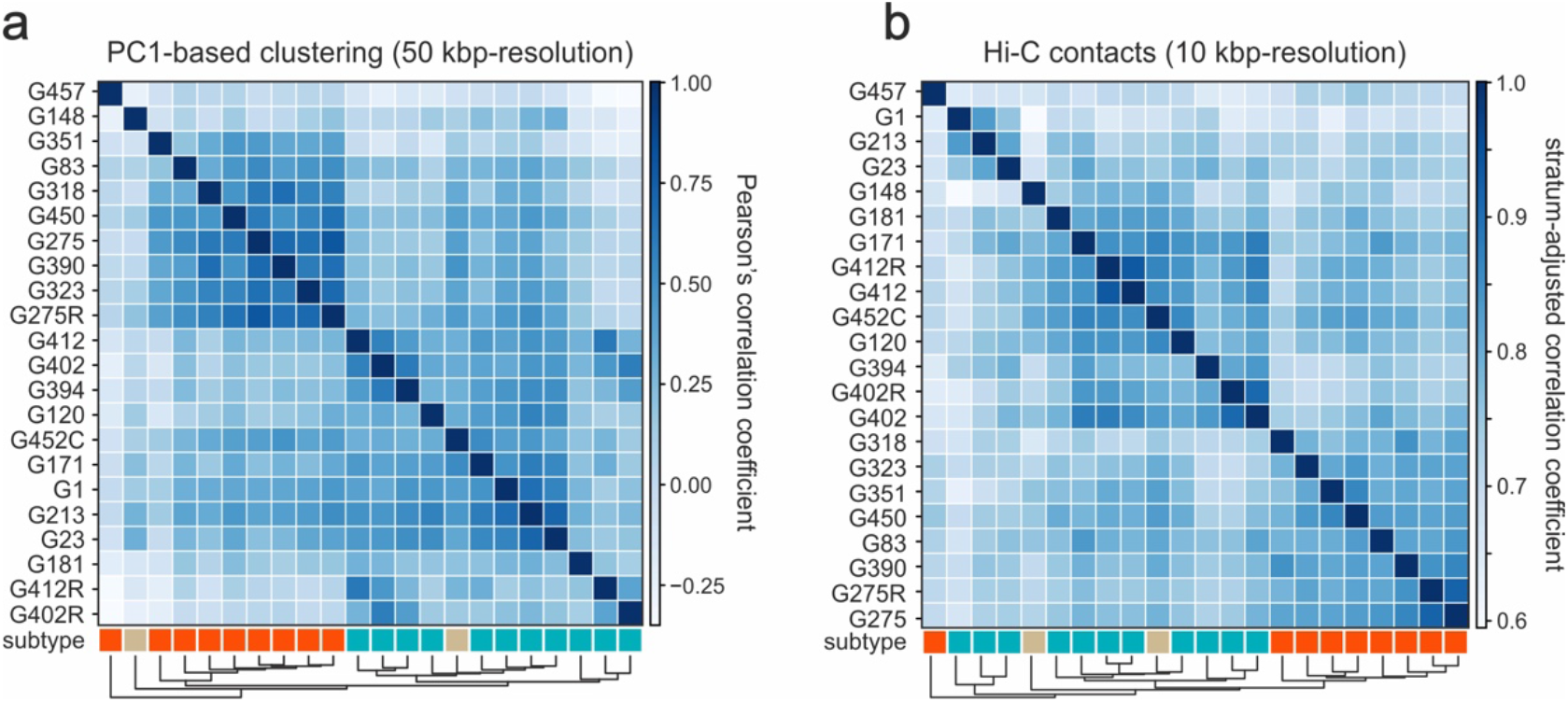
Discriminating GSC subtypes based on 3D genome organization features. **a**, Heatmap showing correlation and unsupervised clustering on the basis of PC1 values called at 50-kbp resolution Hi-C data for 22 GSc lines. The color code (below) reflects the subtype of each line (proneural – brown; mesenchymal – orange; classical – green). **b**, As in panel a, but computing SCC correlation for all Hi-C contacts at 10-kbp resolution

**Supplementary Fig. 5.**
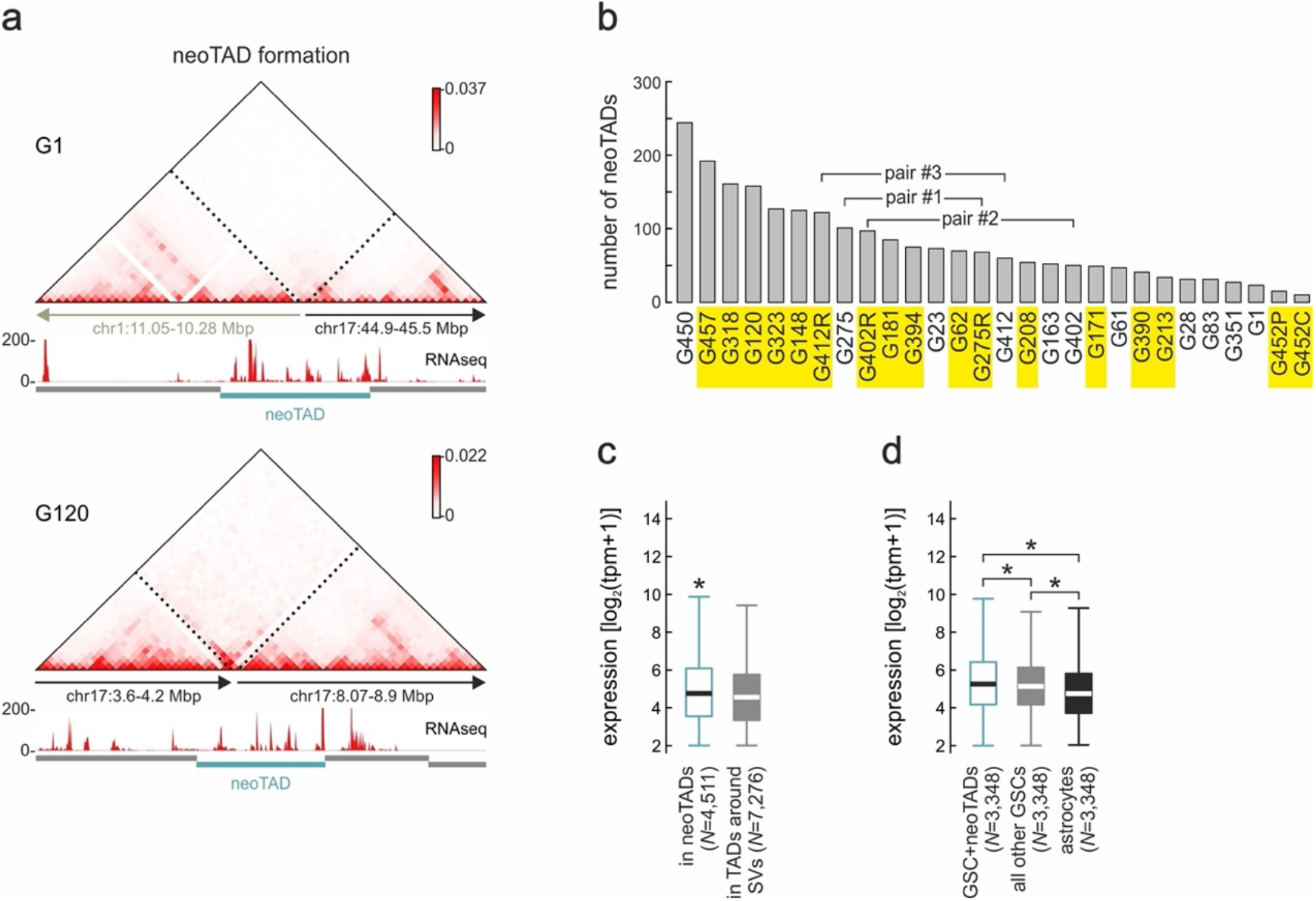
SVs give rise to GSC-specific neoTADs. **a**, Exemplary Hi-C contact maps from G1 and G120 in the 1-1.5 Mbp around a translocation (top) and deletion breakpoint (bottom) that give rise to neoTADs (green rectangle). **b**, Bar plot showing the number of neoTADs identified in Hi-C data from each GSC line. Lines derived from relapse tumors are indicated (yellow). **c**, Box plots showing mean gene expression in neoTADs (green) versus neighboring TADs (grey). \**P*<0.01, two-sided Mann-Whitney U-test. **d**, As in panel d, but for mean expression of genes in GSC-specific neoTADs (green) versus that in GSCs without neoTADs (grey) or in astrocytes (black). \**P* <0.01, two-sided Wilcoxon rank-sum test.

**Supplementary Fig. 6.**
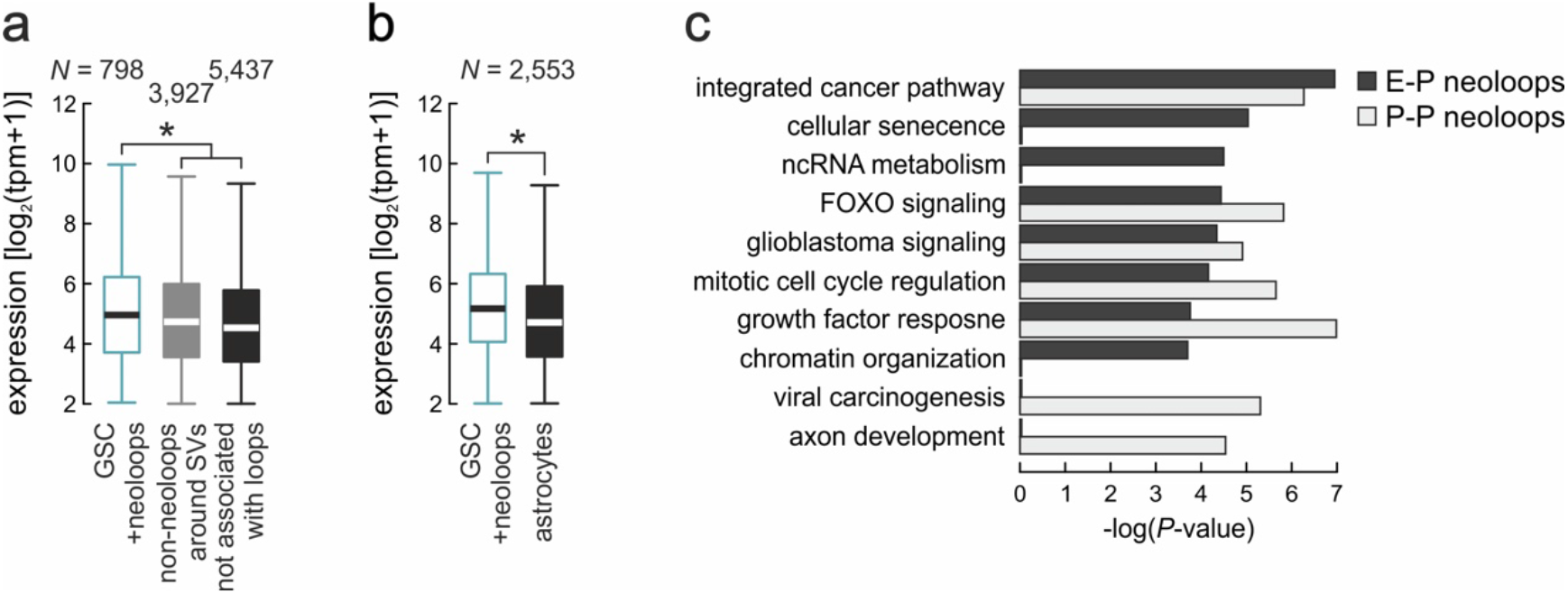
Neoloops sustain higher gene expression in GSCs. **a**, Box plots showing expression of genes associated with neoloops (green) or non-neoloops in GSCs (grey) or not associated with loops (black). *: *P*<0.01, two-sided Mann-Whitney U-test. **b**, As in panel a, but showing mean expression of neoloop-associated genes in GSCs (green) versus the same genes in astrocytes (black). *: *P*<0.01, two-sided Wilcoxon rank-sum test. **c**, GO terms associated with genes connected via E-P (black) or P-P neoloops (light grey).

**Supplementary Fig. 7.**
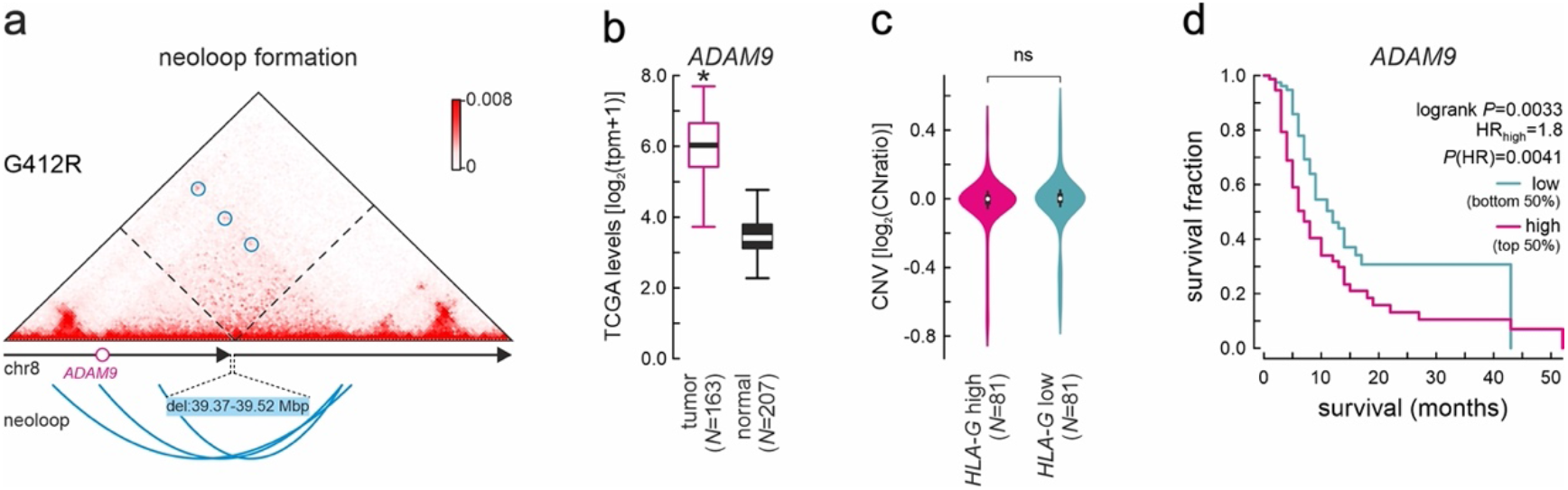
Neoloops in the *ADAM9* locus are associated with poor prognosis. **a**, Exemplary Hi-C contact maps from G412R around a 150-kbp deletion in the *ADAM9* locus. A neoloop forming across the breakpoint is indicated (blue circles). **b**, Box plots showing *ADAM9* expression in TCGA GBM tumor and normal tissue data. *: *P*<0.01, two-sided Mann-Whitney U-test. **c**, Violin plots showing no copy number variation in the *ADAM9* locus from TCGA GBM tumors with high (top 50%, magenta) or low *HLA-G* expression (bottom 50%, green). *: *P*>0.5, two-sided Mann-Whitney U-test. **d**, Kaplan-Meier survival analysis of GBM patients with *ADAM9* high and low expression. *P* - values were calculated using a two-sided log-rank test.

**Supplementary Fig. 8.**
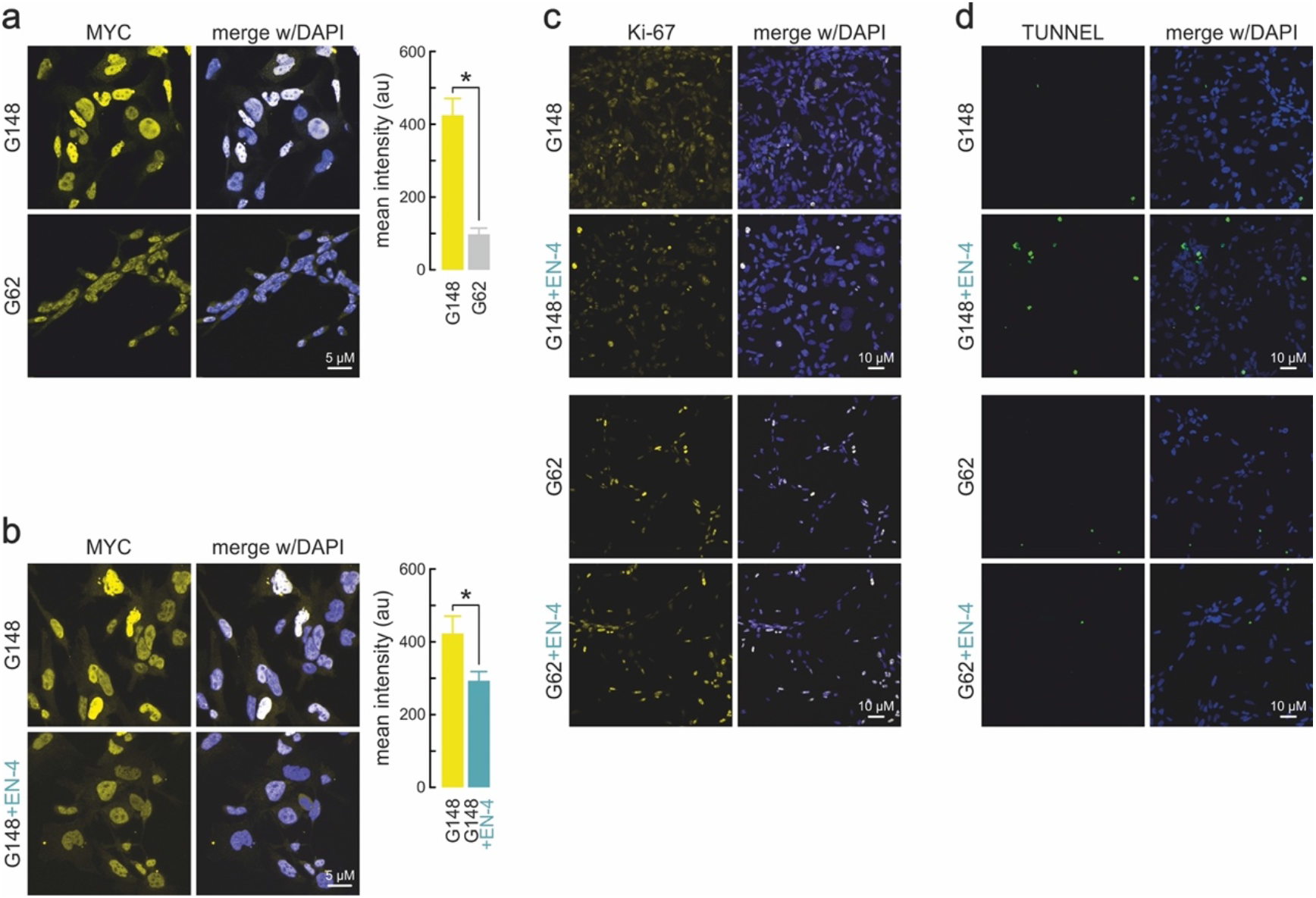
Selective inhibition of MYC-high GSCs by EN-4 treatment. **a**, Left: Representative immunofluorescence images of G148 and G62 cells stained for MYC. Right: Bar plots showing mean MYC levels (±S.D.) in each GSC line. \**P*< 0.01, unpaired two-tailed Student’s t-test. **b**, As in panel a, but G148 stained for MYC after treatment or not with 50 μM EN-4 for 48 h. **c**, As in panel a, but stained for Ki-67 after treatment or not with 50 μM EN-4 for 48 h. **d**, As in panel a, but Tunnel-stained after treatment or not with 50 μM EN-4 for 48 h.

**Supplementary Fig. 9.**
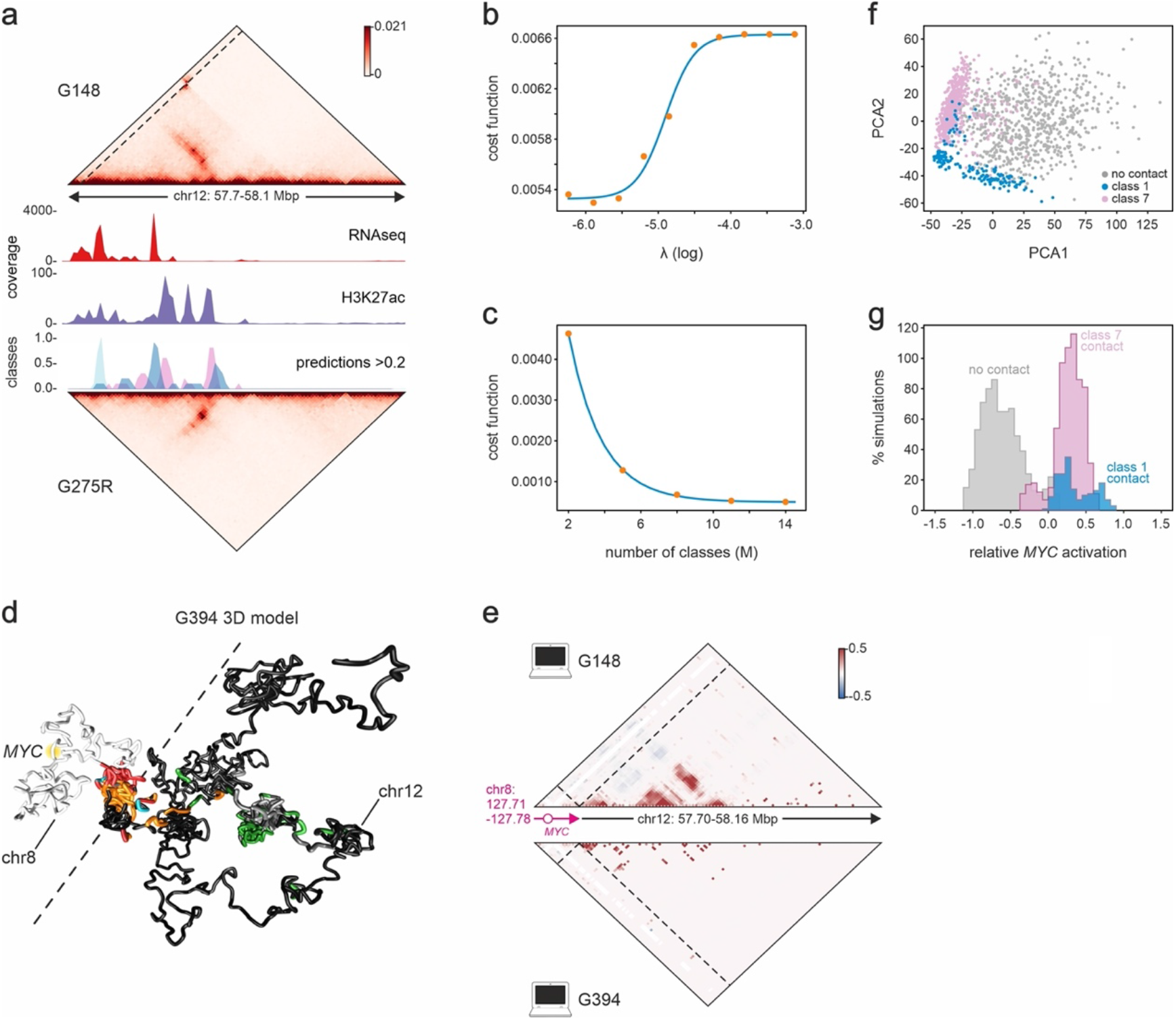
Simulations of the G148-specific chr8:chr12 translocation. **a**, Hi-C contact map of thr 1-Mbp chr12segment of G148 that harbors a translocation with chr8 (dashed line: breakpoint) aligned to classes of polymer beads deduced from simulation in the same G275R region (bottom), and to H3K27ac (middle) and RNA-seq signal tracks (top). **b**, Plot showing the sigmoidal behavior of the cost function with varying regularization constants (λ) that penalize the abundance of binding sites during annealing optimization. **c**, As in panel **b**, but showing exponential decay of the cost function with increasing number of binding site classes (M) during annealing optimization. **d**, Representative 3D rendering of the chr8 (white)-chr12 (black) translocation including *MYC* (yellow halo). Beads from classes that best predict folding are colored (green, red, and orange). **e**, *Top*: Triplet correlation coefficient of *MYC* with all pairs in the simulated G148 translocation. While RNAseq- and H3K27ac-enriched regions form multiple simultaneous contacts with MYC (positive correlation), these occur rather independently or even exclusively of one another (negative correlation). *Bottom*: As above, but for G394 where no contacts form. **f**, PCA clustering of simulated single-allele *MYC* distance profiles (*N*=1500) in G148. Individual models are stratified by the degree of expression and by whether this is due to contacts with beads of enriched H2K27ac (red), RNA-seq (blue) or to lack of contacts (grey). **g**, Per cent of simulated models plotted relative to the extrapolated mean *MYC* activation due to contacts with beads of enriched H2K27ac (red), RNA-seq (blue) or to lack of contacts (grey).

**Supplementary Fig. 10.**
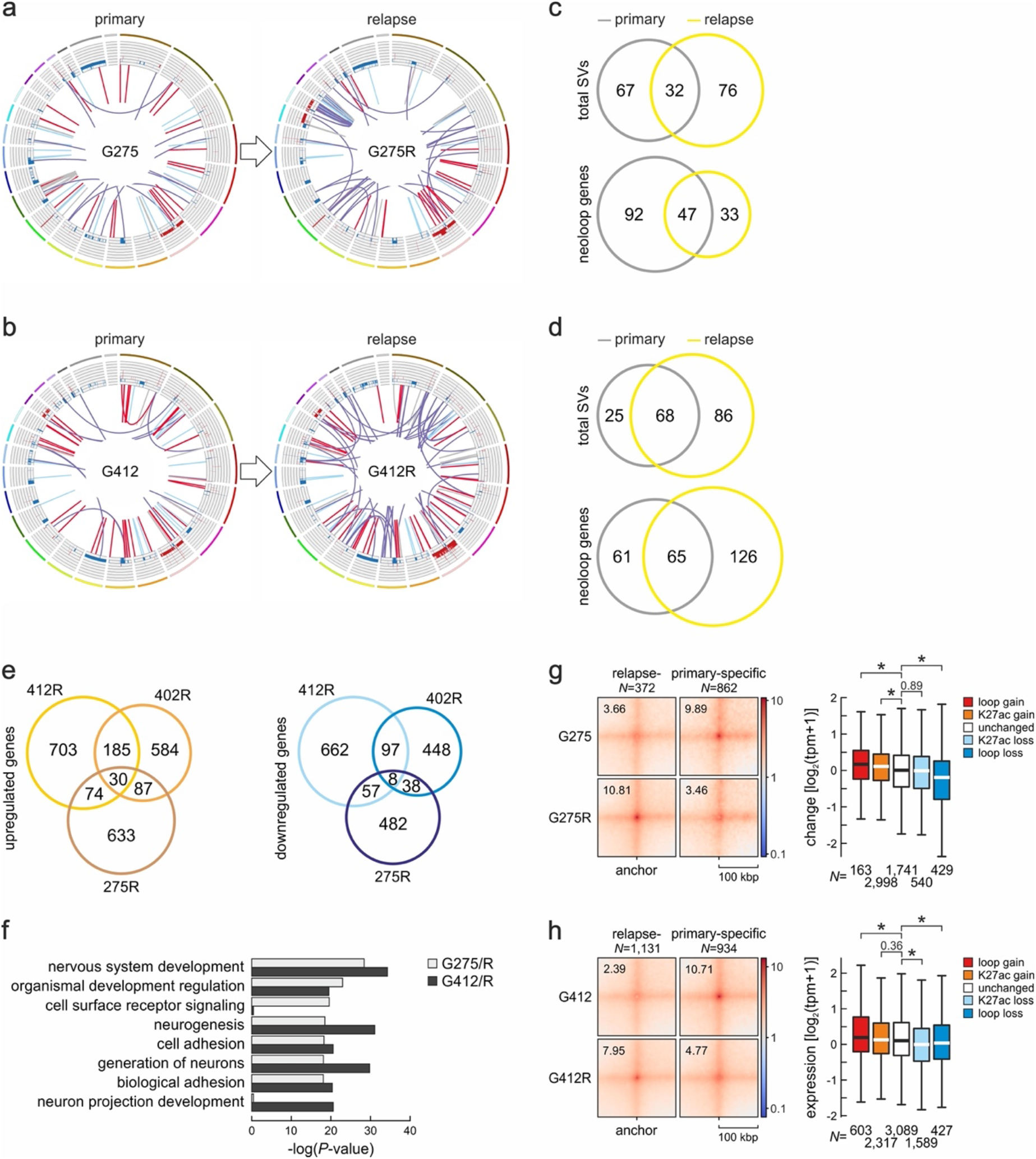
Comparison of SVs in primary versus relapse tumor GSCs. **a**, Circos plot of SVs and CNVs in the G275/275Rprimary-relapse pair. Outer tracks represent chromosomes, inner tracks indicate gain (red: >2 copies) or loss of genomic segments (blue: <2 copies), and lines depict inversions (grey), deletions (light blue), translocations (purple) or duplications (red). **b**, As in panel a, but the G412/412R primary-relapse pair. **c**, Venn diagrams showing shared and unique SVs (top) or neoloop-associated genes (below) in primary (grey) and relapse G275/275R Hi-C data (yellow). **d**, As in panel c, but the G412/412R primary-relapse pair. **e**, Venn diagrams showing shared and unique up- (left) and downregulated genes (right) from all three primary-relapse GSC pairs. **f**, GO terms associated with genes differentially-expressed in the primary versus relapse pairs. **g**, Left: APA plot for all loops specific to the G275 (primary) or G275R (relapse). Right: Box plots showing changes in the expression of genes associated with loops gained (red) or lost (blue), having increased (orange) or decreased H3K27ac (light blue), or not changing upon relapse (white). *: *P*<0.01, two-sided Mann-Whitney U-test. **h**, As in panel g, but for the G412/R pair. *: *P*<0.01, two-sided Mann-Whitney U-test.

**Supplementary Fig. 11.**
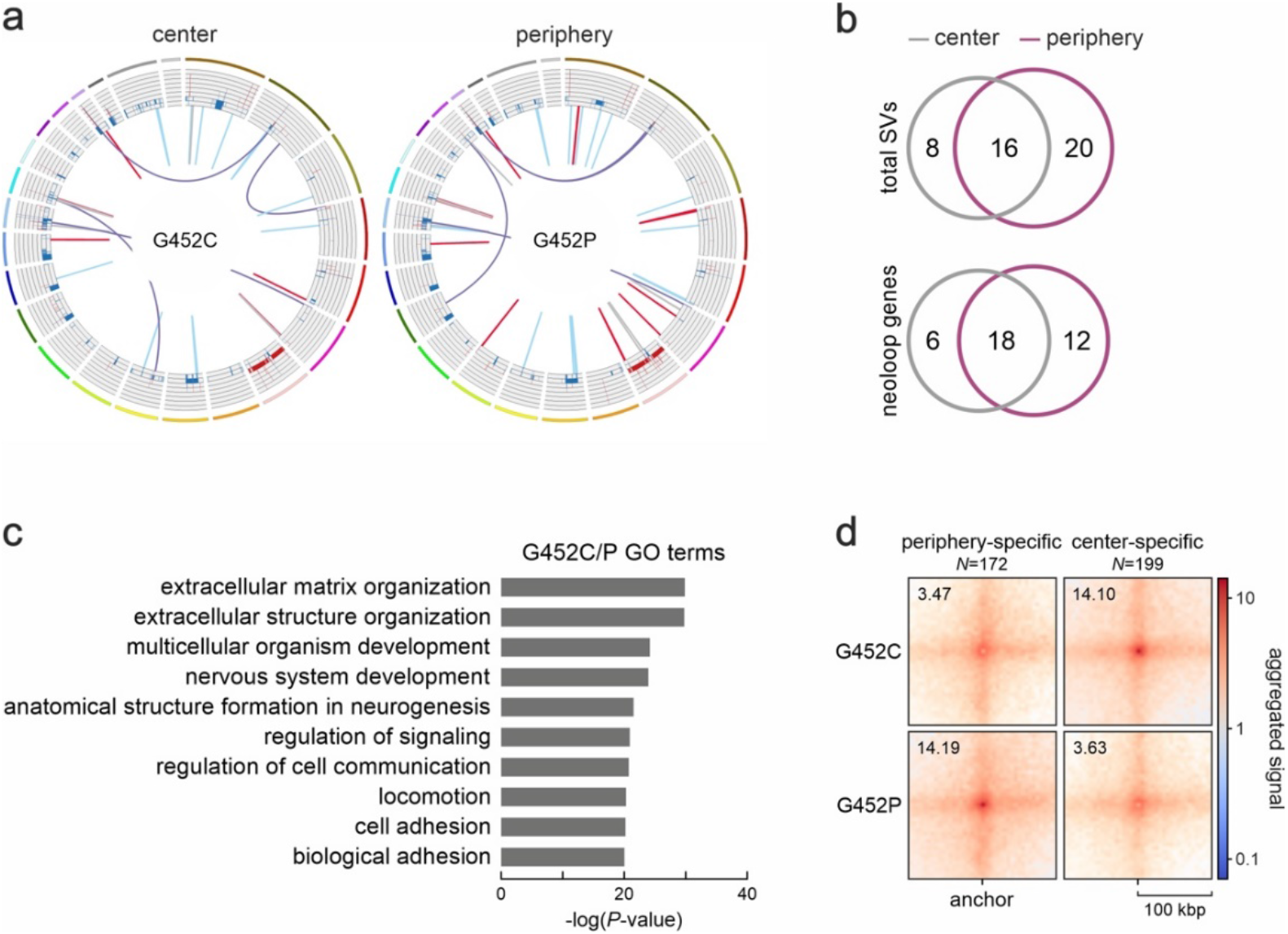
Comparison of SVs in the central versus peripheral part of a GBM tumor. **a**, Circos plot of SVs and CNVs in the G452C/P pair originating from the central and peripheral part of a single GBM tumor. Outer tracks represent chromosomes, inner tracks indicate gain (red: >2 copies) or loss of genomic segments (blue: <2 copies), and lines depict deletions (light blue), inversions (grey), duplications (red) or translocations (purple). **b**, Venn diagrams showing shared and unique SVs (top) or neoloop-associated genes (below) in G452P/C data. **c**, GO terms associated with genes differentially-expressed in the 452C versus the 452P line. **d**, APA plots for all loops specific to the G452C or G452P line.

**Supplementary Table 1.** Clinical features of GBM patients and tumors.

| GSC# | age | sex | sympt | tumor<br>type | tumor<br>location | stupp | MGMT | IDH | EGFR | VEGF | PFS | OS |
| --- | --- | --- | --- | --- | --- | --- | --- | --- | --- | --- | --- | --- |
| 1 | 40 | M | 2.5 | primary | temporal | yes | M | wt | neg | hyper | 6 | 12.5 |
| 23 | 77 | M | 2 | primary | parietal | no | UM | wt | neg | hyper | 1 | 2 |
| 28 | 72 | M | 1.5 | primary | frontal | yes | M | wt | neg | hyper | 6 | 11.5 |
| 61 | 59 | M | 2 | primary | occipital | no | UM | wt | pos | normal | 3 | 6 |
| 62 | 64 | M | 36 | relapse | frontal | yes | M | wt | neg | hyper | 10 | 14 |
| 83 | 52 | M | 0.5 | primary | temporal | yes | UM | wt | pos | hyper | 4 | 8 |
| 120 | 53 | M | 25 | relapse | parietal | yes | UM | wt | neg | hyper | 8 | 16.5 |
| 148 | 55 | M | 6 | relapse | parietal | yes | UM | wt | neg | hyper | 5 | 8 |
| 163 | 56 | M | 5 | primary | parietal | no | UM | wt | neg | normal | 1 | 2 |
| 171 | 74 | M | 13 | relapse | frontal | yes | M | wt | pos | normal | 10 | 17 |
| 181 | 64 | F | 15 | relapse | occipital | yes | M | wt | pos | hyper | 12 | 17 |
| 208 | 66 | M | 22 | relapse | temporal | yes | UM | wt | neg | hyper | 20 | 33 |
| 213 | 50 | M | 18 | relapse | frontal | yes | M | wt | neg | hyper | 9 | 10.5 |
| 275<br>275R | 58 | M | 2 | primary<br>relapse | occipital | yes | M | wt | neg | hyper | 6 | 12 |
| 318 | 69 | M | 25 | relapse | temporal | yes | M | wt | neg | N/A | 21 | 38 |
| 323 | 51 | F | 19 | relapse | parietal | yes | UM | wt | neg | hyper | 4 | 28 |
| 351 | 52 | F | 1 | primary | temporal | yes | M | wt | pos | normal | 84 | 84 |
| 390 | 49 | M | 0.5 | relapse | temporal | yes | M | wt | pos | hyper | 23 | 32.5 |
| 394 | 64 | M | 1 | relapse | frontal | yes | M | wt | pos | hyper | 5 | 23 |
| 402<br>402R* | 58 | M | 0.5 | primary<br>relapse | parietal | yes | UM | wt | neg | hyper | 9 | 23 |
| 412<br>412R** | 56 | F | 1 | primary<br>relapse | frontal | no | UM | wt | neg | hyper | 22 | 31 |
| 450 | 76 | F | 1 | primary | temporal | no | M | wt | neg | hyper | 3 | 6 |
| 452P/C<sup>§</sup> | 67 | F | 46 | relapse | temporal | no | M | wt | pos | hyper | 1.5 | 2 |
| 457 | 59 | F | 0.5 | relapse | frontal | yes | M | wt | pos | hyper | 9.5 | 14.5 |

Age is displayed in years; symptoms' duration is displayed in months; M/UM, methylated/unmethylated; N/A, not available; PFS, progression-free survival displayed in months; OS, overall survival displayed in months; \*initially named 428; \*\*initially named 486; <sup>§</sup>P/C, biopsy peripheral/central to the tumor.

**Supplementary Table 2.**
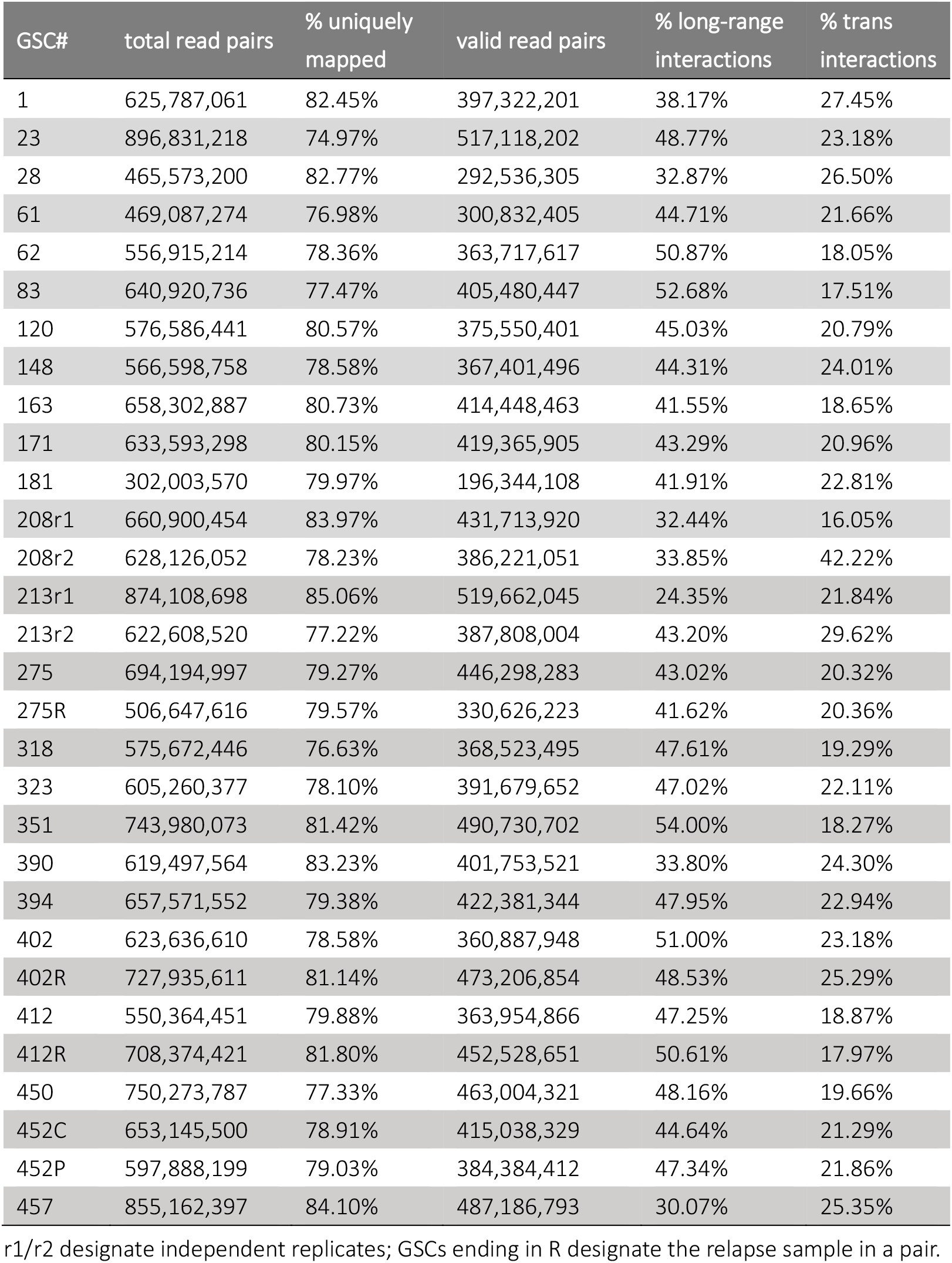
Statistics and quality metrics of Hi-C experiments.

| GSC# | total read pairs | % uniquely mapped | valid read pairs | % long-range interactions | % trans interactions |
| --- | --- | --- | --- | --- | --- |
| 1 | 625,787,061 | 82.45% | 397,322,201 | 38.17% | 27.45% |
| 23 | 896,831,218 | 74.97% | 517,118,202 | 48.77% | 23.18% |
| 28 | 465,573,200 | 82.77% | 292,536,305 | 32.87% | 26.50% |
| 61 | 469,087,274 | 76.98% | 300,832,405 | 44.71% | 21.66% |
| 62 | 556,915,214 | 78.36% | 363,717,617 | 50.87% | 18.05% |
| 83 | 640,920,736 | 77.47% | 405,480,447 | 52.68% | 17.51% |
| 120 | 576,586,441 | 80.57% | 375,550,401 | 45.03% | 20.79% |
| 148 | 566,598,758 | 78.58% | 367,401,496 | 44.31% | 24.01% |
| 163 | 658,302,887 | 80.73% | 414,448,463 | 41.55% | 18.65% |
| 171 | 633,593,298 | 80.15% | 419,365,905 | 43.29% | 20.96% |
| 181 | 302,003,570 | 79.97% | 196,344,108 | 41.91% | 22.81% |
| 208r1 | 660,900,454 | 83.97% | 431,713,920 | 32.44% | 16.05% |
| 208r2 | 628,126,052 | 78.23% | 386,221,051 | 33.85% | 42.22% |
| 213r1 | 874,108,698 | 85.06% | 519,662,045 | 24.35% | 21.84% |
| 213r2 | 622,608,520 | 77.22% | 387,808,004 | 43.20% | 29.62% |
| 275 | 694,194,997 | 79.27% | 446,298,283 | 43.02% | 20.32% |
| 275R | 506,647,616 | 79.57% | 330,626,223 | 41.62% | 20.36% |
| 318 | 575,672,446 | 76.63% | 368,523,495 | 47.61% | 19.29% |
| 323 | 605,260,377 | 78.10% | 391,679,652 | 47.02% | 22.11% |
| 351 | 743,980,073 | 81.42% | 490,730,702 | 54.00% | 18.27% |
| 390 | 619,497,564 | 83.23% | 401,753,521 | 33.80% | 24.30% |
| 394 | 657,571,552 | 79.38% | 422,381,344 | 47.95% | 22.94% |
| 402 | 623,636,610 | 78.58% | 360,887,948 | 51.00% | 23.18% |
| 402R | 727,935,611 | 81.14% | 473,206,854 | 48.53% | 25.29% |
| 412 | 550,364,451 | 79.88% | 363,954,866 | 47.25% | 18.87% |
| 412R | 708,374,421 | 81.80% | 452,528,651 | 50.61% | 17.97% |
| 450 | 750,273,787 | 77.33% | 463,004,321 | 48.16% | 19.66% |
| 452C | 653,145,500 | 78.91% | 415,038,329 | 44.64% | 21.29% |
| 452P | 597,888,199 | 79.03% | 384,384,412 | 47.34% | 21.86% |
| 457 | 855,162,397 | 84.10% | 487,186,793 | 30.07% | 25.35% |

r1/r2 designate independent replicates; GSCs ending in R designate the relapse sample in a pair.

**Supplementary Table 3.** Catalogue of all CNVs, SVs, neoloops, and gene fusion events identified using Hi-C in all GSCs (provided in .xlsx format).

## Notes

### Competing Interest Statement

The authors have declared no competing interest.

